# Dectin-1 limits central nervous system autoimmunity through a non-canonical pathway

**DOI:** 10.1101/2020.05.06.080481

**Authors:** M. Elizabeth Deerhake, Keiko Danzaki, Makoto Inoue, Emre D. Cardakli, Toshiaki Nonaka, Nupur Aggarwal, William E. Barclay, Ru Rong Ji, Mari L. Shinohara

## Abstract

Pathologic roles for innate immunity in neurologic disorders are well-described, but protective aspects of the immune response are less understood. Dectin-1, a C-type lectin receptor (CLR), is largely known to induce inflammation. However, we found that Dectin-1 is protective in experimental autoimmune encephalomyelitis (EAE), while its canonical signaling mediator, Card9, promotes the disease. Notably, Dectin-1 does not respond to heat-killed *Mycobacteria*, an adjuvant to induce EAE. Myeloid cells mediate the protective function of Dectin-1 in EAE and upregulate gene expression of neuroprotective molecules, including Oncostatin M (Osm) through a non-canonical Card9-independent pathway, mediated by NFAT. Furthermore, we found that the Osm receptor (OsmR) functions specifically in astrocytes to reduce EAE severity. Our study revealed a new mechanism of protective myeloid-astrocyte crosstalk regulated by a non-canonical Dectin-1 pathway and identifies novel therapeutic targets for CNS autoimmunity.

**Graphical Abstract:**

- Dectin-1 is a protective C-type lectin receptor (CLR) in experimental autoimmune encephalomyelitis (EAE)
- Dectin-1 promotes expression of *Osm*, a neuroprotective IL-6 family cytokine, in myeloid cells
- OsmR signaling in astrocytes limits EAE progression and promotes remission
- Non-canonical Card9-independent signaling drives a distinct Dectin-1-mediated transcriptional program to induce expression of *Osm* and other factors with protective or anti-inflammatory functions

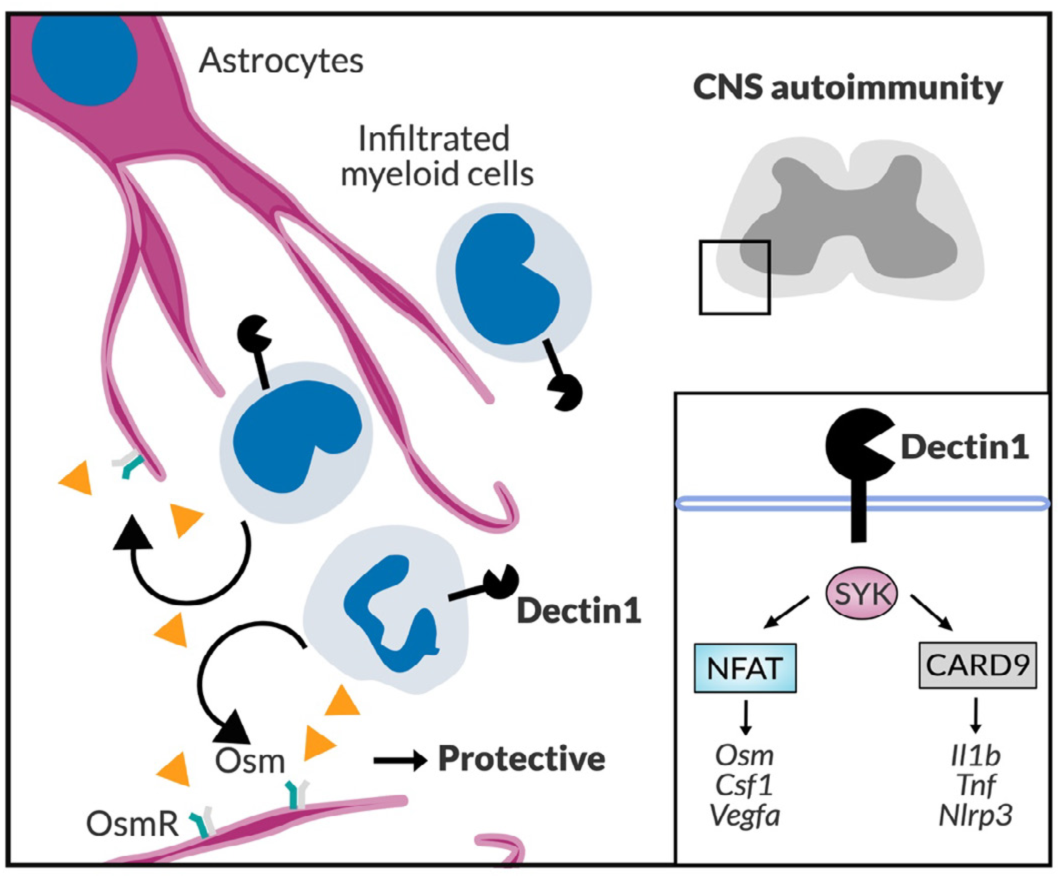

## INTRODUCTION

The innate immune response contributes to both damage and repair in central nervous system (CNS) autoimmunity, and pattern recognition receptors (PRRs) are key to orchestrating innate immune responses^1, 2^. Among PRRs, the C-type lectin receptor (CLR) family, including Dectin-1, has been studied mainly in fungal infections and remains less characterized in sterile inflammation and autoimmune disorders, including multiple sclerosis (MS)^1^. Dectin-1 induces IL-1β and resulting Th17 responses during fungal infections via Card9/NFκB signaling^3, 4^. Although Dectin-1 is known as a receptor for fungal β-glucans^5^, some studies have identified host-derived endogenous ligands for Dectin-1, as well as functions of Dectin-1 signaling beyond the setting of fungal infections^6, 7, 8, 9, 10^. Notably, in one^11^ of three reports^12, 13^ of experimental autoimmune uveitis (EAU), Dectin-1 was found to be detrimental. Dectin-1 also appears to be pathogenic in animal models of spinal cord injury^14^ and stroke^15^. Nevertheless, the function of Dectin-1 in CNS disorders may depend on the type of neuropathology, and the role of Dectin-1 in autoimmunity and neuroinflammation is still largely unexplored.

Given the well-described pro-inflammatory functions of Dectin-1 signaling, one may predict that Dectin-1 would exacerbate neuroinflammation in the EAE model of MS. However, this study demonstrates that Dectin-1 signaling in myeloid cells limits neuroinflammation and is protective in EAE, although a major Dectin-1 signaling molecule, Card9, promotes disease development. We found that non-canonical Card9-independent Dectin-1 signaling involving NFAT drives expression of Oncostatin M (Osm), an IL-6 family cytokine with neuroprotective functions^16, 17, 18^, and signaling through the Osm receptor (OsmR) on astrocytes reduces disease progression and promotes recovery in EAE. Furthermore, we identified a Card9-independent Dectin-1-mediated transcriptional program driving expression of *Osm* and other neuroprotective genes. Our findings provoke a re-consideration of Dectin-1 signaling and functions by identifying a new mechanism of protective myeloid-astrocyte communication in CNS autoimmunity.

## RESULTS

### Elevated gene expression of the C-type lectin receptor, Dectin-1 (*CLEC7A*) in MS lesions

We evaluated gene expression of CLRs in multiple datasets profiling MS brain lesions^18^. Among genes encoding CLRs with known immune functions, *CLEC7A* (Dectin-1) and *CLEC4A* (DCIR), closely linked on chromosome 12, were notable for their elevated expression in MS brain specimens in two independent datasets (Supplementary Fig. 1a-c)^19, 20, 21^, suggesting possible involvement of Dectin-1 and DCIR in MS. In EAE, DCIR has been reported to be protective^22, 23^ but the function of Dectin-1 in CNS autoimmunity remained unknown.

**Figure 1.**
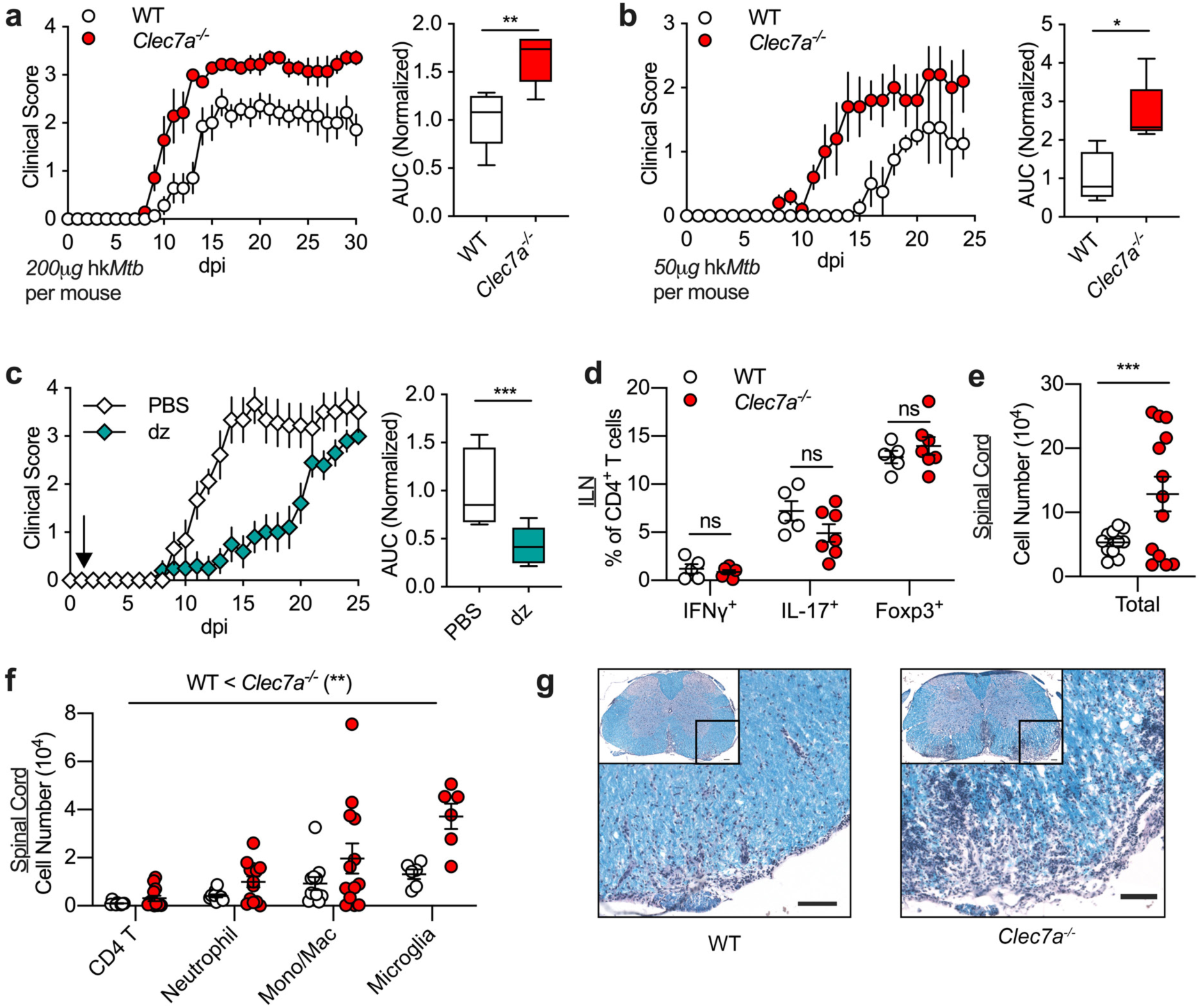
Dectin-1 limits neuroinflammation in EAE. **(a-c)** EAE clinical scores (left panels) and AUC (right panels) for statistical analysis. Comparison between WT and *Clec7a*^*-/-*^ mice using standard immunization (200μg hk*Mtb*/mouse) with *n*=7 mice/group (a) or reduced adjuvant immunization (100μg hk*Mtb*/mouse) with *n*=4 WT and *n*=5 *Clec7a*^*-/-*^ mice/group (b). Comparison of WT mice receiving either hot alkali-depleted zymosan (dz) (500μg/mouse) or PBS *i*.*v*. at 1-dpi using *n*=9 PBS and *n*=10 dz-treated mice/group using standard immunization (c). Data representative of >3 independent experiments (a) or 2 independent experiments each (b, c). **(d-f)** Frequency of Th cell subsets in ILN between WT and *Clec7a*^*-/-*^ mice at 9-dpi (d). Data representative of 3 independent experiments. Total cell numbers (e) and numbers of indicated cell types (f) in spinal cord. Significant main effect of genotype (WT < *Clec7a*^*-/-*^) (**) by 2-factor RM-ANOVA indicated in (f). Data are combined from three independent experiments. One datapoint denotes a result from one mouse. **(g)** LFB-PAS staining of lumbar SC in WT and *Clec7a*^*-/-*^ mice at EAE 17-dpi. Scale 100μm. Representative images from 10 mice/group combined from two independent experiments.

### Dectin-1 is a protective C-type lectin receptor in EAE

Next, we sought to test the function of Dectin-1 in CNS autoimmunity using EAE induced with the MOG_35−55_ autoantigen peptide. We initially hypothesized that Dectin-1 may exacerbate EAE severity by promoting IL-1β expression and Th17 differentiation, given its known role in antifungal immunity^3, 4^ and the pathogenicity of Th17 cells in EAE development^24^. Instead, we found that Dectin-1-deficient (*Clec7a*^*-/-*^) mice developed more severe disease than WT mice (Fig. 1a). A mild EAE induction with a reduced amount of adjuvant also showed the protective effect of Dectin-1 (Fig. 1b). Next, we tested whether administering a Dectin-1 agonist was sufficient to limit EAE severity. To test this, we used hot alkali-depleted zymosan (d-zymosan), which is specific to Dectin-1 but does not stimulate TLR2^25^. When d-zymosan was subcutaneously administered together with heat-killed *Mtb* (hkMtb) adjuvant and antigen, we did not observe any difference in EAE severity (Supplementary Fig. 2a). However, a single intravenous (*i*.*v*.) injection of d-zymosan on one day post-immunization (dpi) significantly inhibited EAE development (Fig. 1c). Together, these results indicate that Dectin-1 limits EAE severity, likely acting beyond the site of immunization.

**Figure 2.**
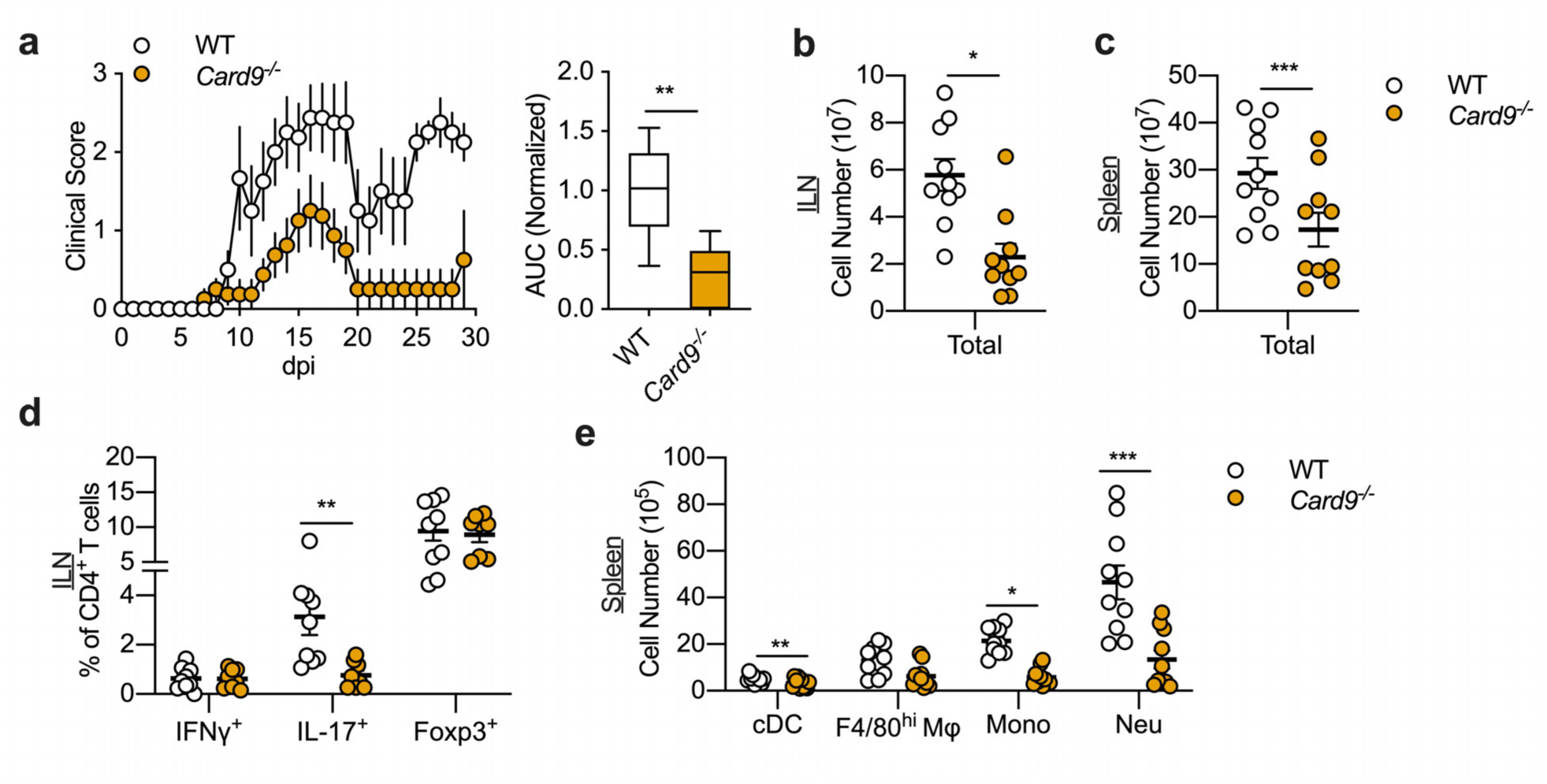
Card9 exacerbates EAE and promotes a Th17 response. **(a)** EAE clinical scores (left panel) and AUC (right panel) for statistical analysis. Comparison between WT (*n*=9) and *Card9*^*-/-*^ (*n*=8) mice. Data are combined from two independent experiments. **(b, c)** Total cell numbers in ILN (b) and spleen (c) between WT and *Card9*^*-/-*^ mice at 9-dpi. **(d, e)** Frequency of Th cell subsets in ILN (d) and numbers of indicated cell types in spleen (e). One datapoint denotes a result from one mouse. Data are combined from three independent experiments.

### Dectin-1 limits CNS inflammation in EAE

To determine how Dectin-1 regulates EAE, we evaluated immune cell subsets by flow cytometry in secondary lymphoid organs and in the spinal cord (SC) of WT and *Clec7a*^*-/-*^ mice. Unexpectedly, numbers of T_H_1, T_H_17, or T_reg_ cells in the draining inguinal lymph nodes (ILN) (Fig. 1d, Supplementary Fig. 2b) were similar between WT and *Clec7a*^*-/-*^ mice around the time of EAE onset (9-dpi), suggesting no significant effect on CD4^+^ T cell polarization. Other myeloid and lymphocyte cell types also showed similar numbers between WT and *Clec7a*^*-/-*^ mice in spleen and LN at 9-dpi (Supplementary Fig. 2c, d). In contrast, *Clec7a*^*-/-*^ mice had elevated immune cell infiltration in the SC at 9-dpi compared to WT controls (Fig. 1e) across multiple immune cell subsets (Fig. 1f), suggesting that Dectin-1 broadly limits inflammation in the SC during EAE. We also observed increased demyelination in SCs of *Clec7a*^*-/-*^ mice by Luxol fast blue/periodic acid-Schiff (LFB-PAS) staining (Fig. 1g; Supplementary Fig. 2e). Together, this demonstrates that Dectin-1 specifically limits CNS inflammation during EAE.

To test whether Dectin-1 reduces immune cell migration to the CNS in a cell-intrinsic manner, we generated mixed bone-marrow (BM) chimeras reconstituted with both WT and *Clec7a*^*-/-*^ BM cells (Supplementary Fig. 2f, g). Following EAE induction, the relative proportion of CD11b^+^ myeloid cells derived from WT and *Clec7a*^*-/-*^ BM cells were comparable in the spleen and the SC (Supplementary Fig. 2h), suggesting Dectin-1 does not limit myeloid cell infiltration into the CNS in a cell-intrinsic manner. In summary, Dectin-1 decreases neuroinflammation in EAE, but this effect was not attributable to altered T helper cell polarization or cell-intrinsic defects in myeloid cell migration into the CNS.

### Card9 exacerbates EAE and promotes Th17 responses

We next evaluated the role of Card9, a major mediator of Dectin-1 signaling^3^, in EAE. Remarkably, *Card9*^*-/-*^ mice were resistant to EAE (Fig. 2a), indicating that Card9, unlike Dectin-1, is pathogenic in EAE. The immune cell profile of *Card9*^*-/-*^ mice in EAE is also clearly distinct from that of *Clec7a*^*-/-*^ mice: Total cell numbers in spleen and ILNs were significantly reduced in *Card9*^*-/-*^ mice at 9-dpi (Fig. 2b, c) with specific reductions of T_H_17 cells and macrophages in ILNs (Fig. 2d; Supplementary Fig. 3a) and cDCs, monocytes, and neutrophils in the spleen (Fig. 2e; Supplementary Fig. 3b). Next, we used a BM chimera system to test whether immune cells were sufficient to mediate Card9 function in EAE. Indeed, recipients reconstituted with *Card9*^*-/-*^ BM cells were resistant to EAE development, indicating that BM-derived immune cells are responsible for Card9 function (Supplementary Fig. 3g). In summary, we found that although Dectin-1 is protective in EAE, Card9 is pathogenic.

**Figure 3.**
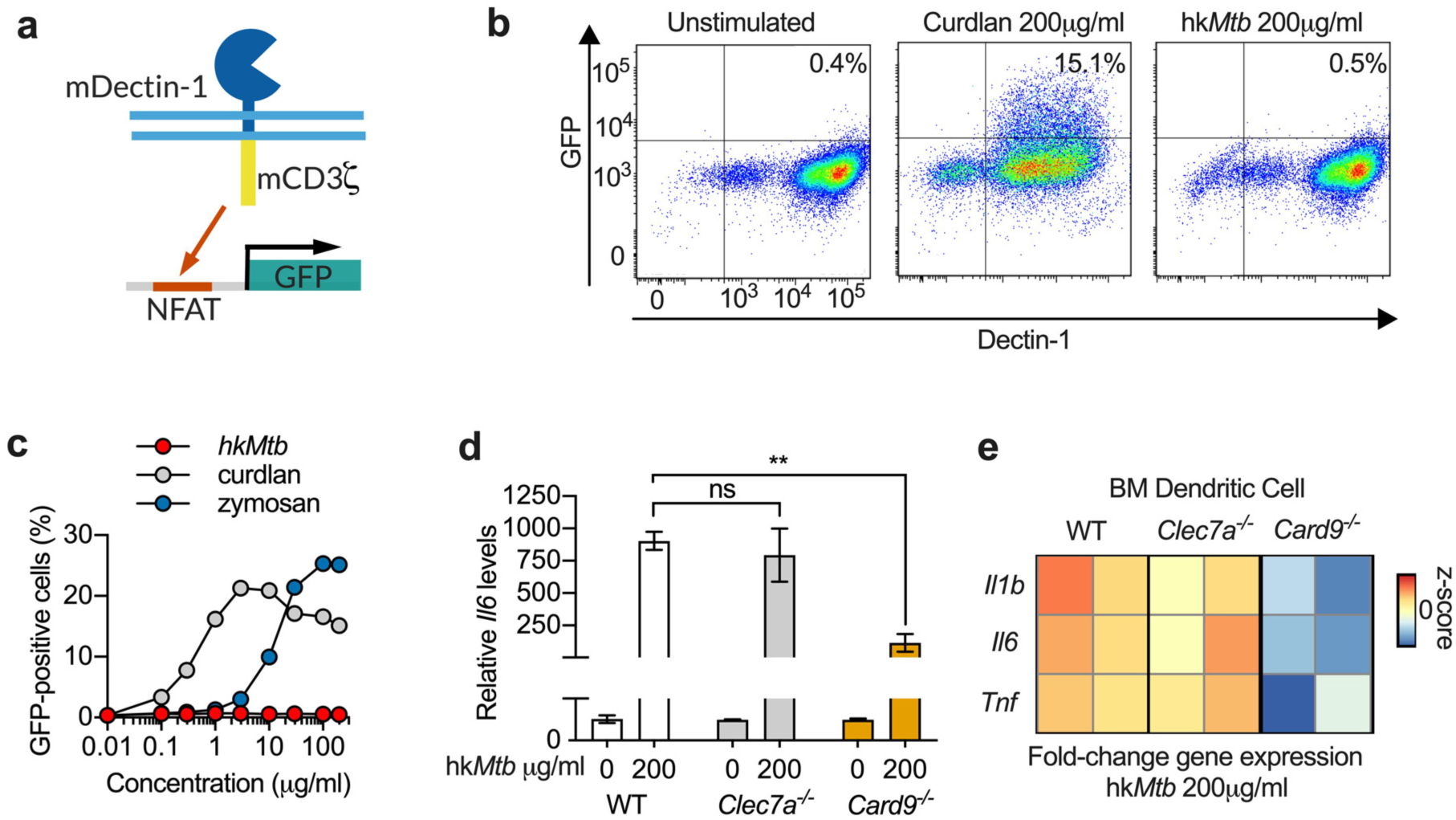
Dectin-1 does not respond to hk*Mtb*, EAE adjuvant. **(a)** Schematic of the mDectin-1/mCD3ζ-GFP reporter cell line. Extracellular and trans-membrane domains of mouse Dectin-1 were fused to the cytoplasmic domain of CD3ζ. Activation of Dectin-1 was detected by a GFP reporter. **(b)** Representative flow cytometry results, showing Dectin-1-GFP reporter at 20-hrs with or without curdlan (200 μg/ml) or hk*Mtb* (200 μg/ml) stimulation in tissue culture. Values indicate percentages of GFP^+^Dectin-1^+^ cells. **(c)** Proportions of GFP^+^Dectin1^+^ mDectin-1/mCD3ζ-GFP reporter cells with titrated concentrations of hk*Mtb*, zymosan, or curdlan with indicated concentrations. **(d, e)** RT-qPCR analysis of *Il6* normalized to unstimulated control (d) and heatmap of *Il1b, Il6*, and *Tnf* expression normalized to unstimulated control (e), showing z-scores. WT, *Clec7a*^*-/-*^, and *Card9*^*-/-*^ BMDCs stimulated with 200 μg/ml hk*Mtb* in tissue culture for 3 hrs. Mean ± SEM of duplicate wells shown, with data representative of two independent experiments.

### Impact of Dectin-1 on EAE is not attributed to hk*Mtb*, used in EAE induction

Next, we sought to determine whether Dectin-1 function in EAE is attributed to the disease induction method which uses hk*Mtb* as an adjuvant. To evaluate whether Dectin-1 recognizes hk*Mtb*, we generated a Dectin-1 signaling reporter cell line, in which a chimeric protein (consisting of mouse Dectin-1 extracellular and transmembrane domains fused to a mouse CD3ζ cytoplasmic domain^26^) is expressed in a GFP reporter T cell hybridoma^27^ (Fig. 3a). Although zymosan and a Dectin-1-specific ligand, curdlan, induced robust GFP reporter expression, hk*Mtb* did not induce GFP expression, even at high concentrations (Fig. 3b, c). Control cells, transfected with a plasmid lacking the chimeric Dectin-1 gene, showed no response to either stimulation, confirming the specificity of the Dectin-1 reporter system (Supplementary Fig. 3h).

To confirm that Dectin-1 does not respond to hk*Mtb* with primary cells, we cultured BM-derived dendritic cells (BMDCs) with hk*Mtb* and evaluated pro-inflammatory cytokine expression. Indeed, *Il1b, Il6*, and *Tnf* mRNA expression was not altered in the absence of Dectin-1 (Fig. 3d, e). In contrast, the absence of Card9 significantly reduced upregulation of *Il1b, Il6*, and *Tnf* mRNA expression upon hk*Mtb* stimulation (Fig. 3d, e). Thus, while Dectin-1 does not detect hk*Mtb*, Card9 mediates response to adjuvant likely through other receptors upstream of Card9, as previously reported^13, 28^. These results indicate that the protective function of Dectin-1 is not attributable to recognition of hk*Mtb*. Together, these findings argue for the involvement of other stimuli, including endogenous Dectin-1 ligands^6, 9, 29^, in mediating Dectin-1 function in EAE.

### Peripheral and CNS-resident myeloid cells express Dectin-1 in EAE

We next evaluated Dectin-1 expression under naïve and EAE conditions. Dectin-1 was expressed on neutrophils, monocytes, macrophages, and DCs in the spleen under naïve and EAE conditions, but not on T and B cells (Fig. 4a, Supplementary Fig. 4a). During EAE, neutrophils and monocytes were the predominant Dectin-1-expressing cell types in the spleen (Fig. 4b). In the SC of EAE mice, Dectin-1^+^ cells were seen in white matter by histology, and co-localized with CD11b^+^ signal, indicative of myeloid cells (Fig. 4c). We observed high Dectin-1 expression in a subset of Iba1^+^ macrophages/microglia with amoeboid morphology in EAE lesions, but less Dectin-1 expression on Iba1^+^ cells with a ramified microglia-like morphology (Supplementary Fig. 4b). Although Dectin-1 expression was not detected on Tmem119^+^ microglia^30^ by immunofluorescent histology (Supplementary Fig. 4c), flow-cytometry did detect significant upregulation of Dectin-1 expression in microglia (Tmem119^+^CD11b^lo^CD45^lo^) during EAE (Fig. 4d, e). Despite upregulating Dectin-1, the expression levels of Dectin-1 in microglia were not as high as those of infiltrated myeloid cells in EAE, including neutrophils and monocytes (Supplementary Fig. 4d; Fig. 4d, e).

**Figure 4.**
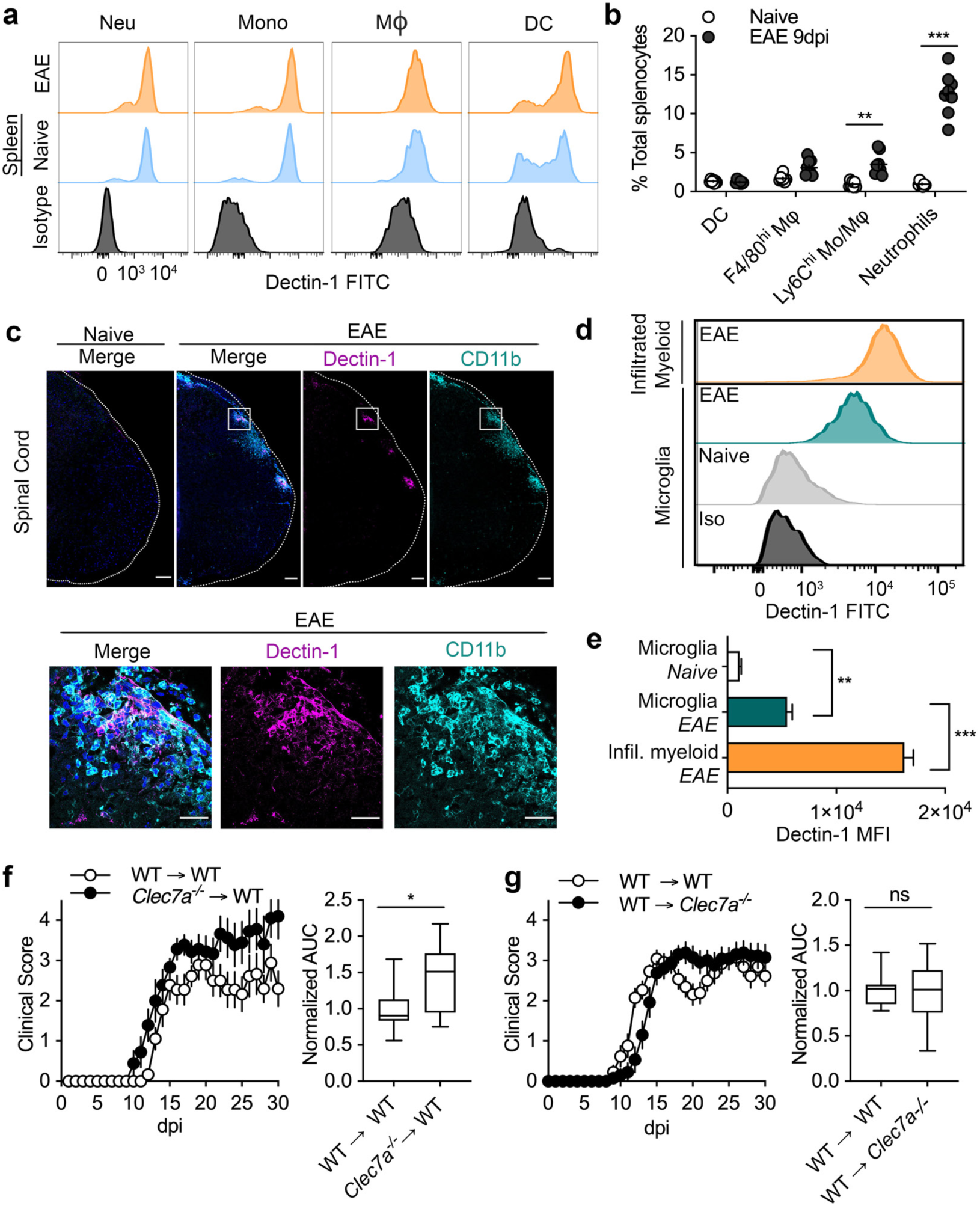
Cell type-specific Dectin-1 expression and function. **(a)** Flow cytometry histograms, indicating Dectin-1 expression in splenic neutrophils (CD11b^+^Ly6G^+^), monocytes (CD11b^+^Ly6C^+^Ly6G^-^), macrophages (CD11b^+^F4/80^hi^), and DCs (CD11b^+^CD11c^+^) from naïve and EAE 9-dpi WT mice compared to isotype control. **(b)** Frequency of myeloid cell subsets in total splenocytes in naïve (*n*=7) and EAE 9-dpi (*n=*8) mice. Data are combined from 2 independent experiments. **(c)** Localization of Dectin-1 expressing cells with CD11b counter-staining in the lumbar SC of naïve and EAE 17-dpi mice. Bottom panels are enlarged images of regions indicated with squares in top panels. Scale 100 μm in upper panels and 25 μm in lower panels. **(d, e)** Flow cytometry histograms (d) and MFI (e) of Dectin-1 expression in microglia (CD11b^lo^CD45^lo^Tmem119^+^) and infiltrated myeloid cells (CD11b^hi^CD45^hi^Tmem119^-^) in SC from naïve and EAE 16-dpi mice. *n*=3 mice/group. Data are representative of 2 independent experiments. **(f, g)** EAE scores (left panels) and AUC for statistical analysis (right panels) of irradiation BM chimeras. WT or *Clec7a*^*-/-*^ BM cells into irradiated WT recipients (*n*=9 for both groups) (f). WT BM into WT or *Clec7a*^*-/-*^ recipients using (*n*=13 for both groups) (g). Data are combined from two independent experiments each.

### Cells of hematopoietic origin mediate Dectin-1 function in EAE

To determine whether hematopoietic-derived or CNS-resident myeloid cells mediate the protective role of Dectin-1 during EAE, we generated BM chimeras by adoptively transferring BM cells from *Clec7a*^*-/-*^ or WT mice to WT recipients (Supplementary Fig. 4e) or by transferring BM cells from WT mice to *Clec7a*^*-/-*^ or WT recipients. Chimeras that received *Clec7a*^*-/-*^ BM cells exhibited more severe disease than chimeras which received WT BM (Fig. 4f). Together with our finding that Dectin-1 is expressed by myeloid cells (Fig. 4a), but not by lymphoid cells (Supplementary Fig. 4a), this study demonstrates that hematopoietic-derived myeloid cells are likely to mediate the protective effect of Dectin-1. In reciprocal experiments, no difference was found between *Clec7a*^*-/-*^ and WT recipients reconstituted with WT BM cells (Fig. 4g), suggesting that radiation-resistant cells, including microglia, were less likely to mediate the protective function of Dectin-1. Although the irradiation BM chimera approach does not fully rule out the involvement of microglia^31^, the results strongly suggest that hematopoietic-derived myeloid cells mediate the protective role of Dectin-1 in EAE.

### Dectin-1 promotes expression of neuroprotective cytokine, Oncostatin M (Osm)

Previous studies suggest that CNS-infiltrating myeloid cells may have protective functions in CNS injury and repair^32, 33^. Remarkably, the administration of Dectin-1/TLR2 ligand, zymosan, was also shown to promote optic nerve regeneration^34^. Thus, we wondered whether the protective role of Dectin-1 could be mediated by neuroprotective factors expressed by CNS-infiltrating myeloid cells. To investigate this, we stimulated splenic CD11b^+^ myeloid cells with the Dectin-1 agonist, curdlan, and screened expression of genes encoding secreted proteins with neuroprotective functions^35^. Among the candidates, curdlan potently upregulated the expression of *Osm*^36^, a multi-functional IL-6 family cytokine with neuroprotective roles in different CNS disorders^16, 17, 37^ (Supplementary Fig. 5a). In addition to CD11b^+^ splenocytes (Supplementary Fig. 5a), *Osm* expression was also upregulated by curdlan in primary neutrophils, monocytes, and BM-derived DCs (BMDCs) but to a lesser extent in BM-derived macrophages (BMDMs) (Fig. 5a, b). Upregulation of Osm protein by curdlan stimulation was confirmed in supernatants of neutrophil cell culture, as well as neutrophil cell lysates (Supplementary Fig. 5b).

**Figure 5.**
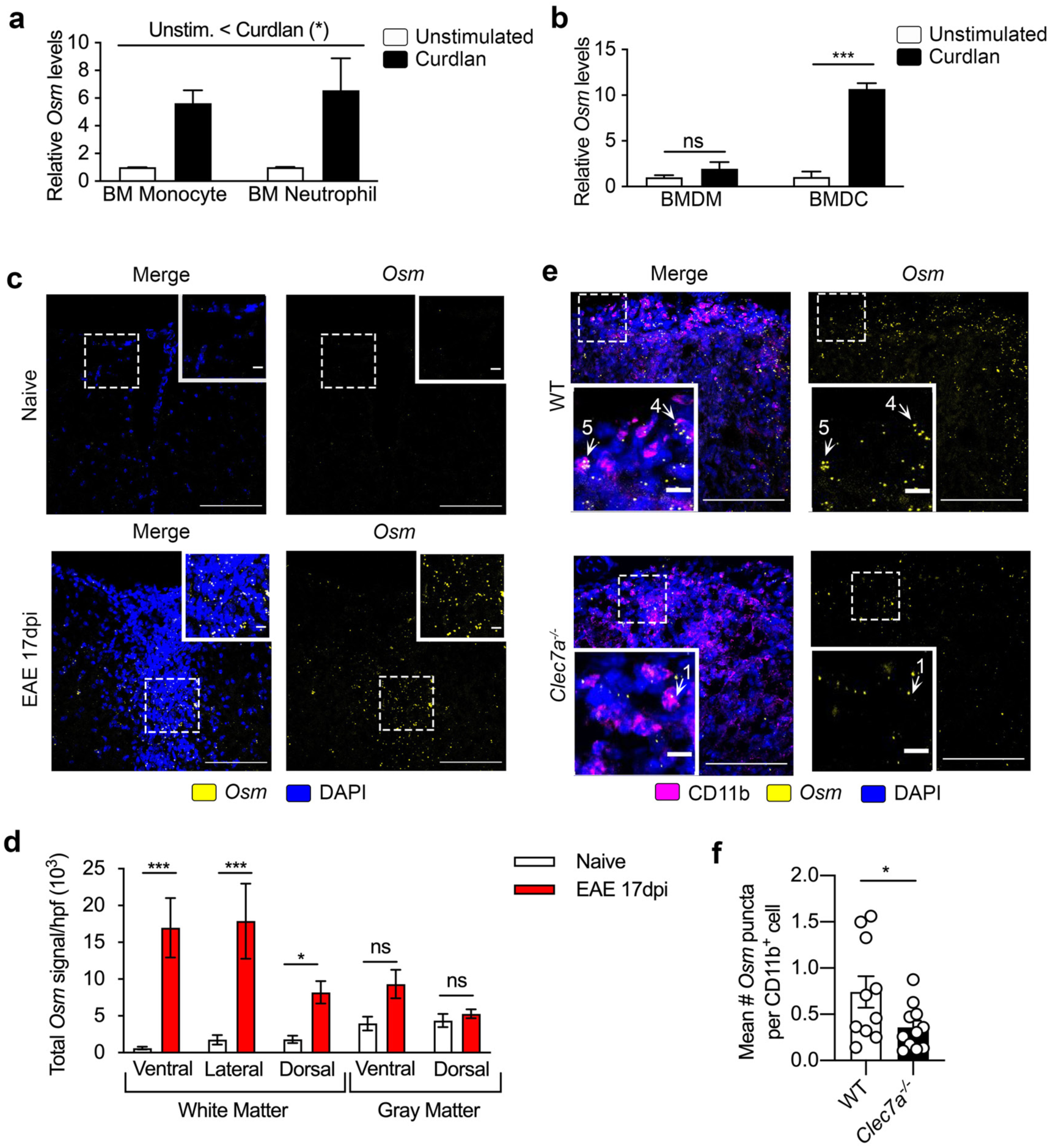
Dectin-1 promotes expression of Oncostatin M (Osm). **(a, b)** *Osm* mRNA levels in indicated myeloid cell types with *ex vivo* curdlan (100 μg/ml) stimulation for 3 hrs. Evaluated are FACS-sorted BM monocytes (CD11b^+^Ly6C^hi^Ly6G^-^) and neutrophils (CD11b^+^Ly6G^+^) (a), as well as GM-CSF-derived BMDCs and M-CSF-derived BMDM (b). Significant main effect of stimulation (Unstimulated < Curdlan) (*) by 2-factor ANOVA indicated in (a). Post-hoc Sidak test following 2-factor ANOVA indicated in (b). Mean ± SEM shown. Data are combined from two independent experiments. **(c)** Representative images of RNAscope *in situ* hybridization (ISH) of *Osm* mRNA in the lumbar SC of WT naïve and EAE 17-dpi mice. **(d)** Region-specific quantification of *Osm* mRNA signal per hpf (sum of integrated density) in lumbar spinal cord regions from WT naïve and EAE 17-dpi mice using *n*=4 mice/group, analyzed by 2-factor ANOVA of log-transformed data with post-hoc Sidak test shown. Representative of two independent experiments. **(e)** Representative images of combined ISH of *Osm* mRNA and CD11b staining in the ventrolateral white matter of lumbar SC from WT and *Clec7a*^*-/-*^ mice at 17-dpi EAE. Scale 100 μm in main panels, 10 μm in inset. **(f)** Quantification of mean *Osm* mRNA puncta per CD11b^+^ cell. One datapoint denotes a result from one mouse. Mean ± SEM. Data from two independent experiments.

We next evaluated whether microglia could also upregulate *Osm* upon curdlan stimulation. Because microglia from naïve mice did not express Dectin-1 (Fig. 4d), we instead isolated Dectin-1-expressing microglia from EAE mice. Specifically, we sorted microglia (CD11b^lo^CD45^lo^) and infiltrating myeloid cells (CD11b^hi^CD45^hi^) from the SC of WT EAE mice and stimulated them with curdlan. While SC-infiltrated myeloid cells showed potent upregulation of *Osm*, microglia had a more muted response despite expressing Dectin-1 (Supplementary Fig. 5c, d). These findings suggest that Dectin-1 signaling drives *Osm* expression in a cell-type-specific manner, whereby microglia and BMDMs show a more limited response than neutrophils, monocytes, and BMDCs.

### Osm expression is upregulated in the CNS during EAE

Osm protein expression is detected in MS lesions by histology^38^ and in culture supernatant of peripheral blood mononuclear cells (PBMCs) from MS patients^39^. In EAE, we found a significant increase in serum Osm protein levels in WT EAE mice at 9-dpi, compared to WT naïve mice (Supplementary Fig. 5e), while serum Osm levels in *Clec7a*^*-/-*^ mice showed a slight but not statistically significant reduction at 9-dpi (Supplementary Fig. 5e). Next, we sought to examine *in situ* Osm expression in the CNS during EAE. Because antibody staining for Osm, a secreted cytokine, was diffuse and challenging to quantify in a cell-specific manner, we used RNAscope *in situ* mRNA hybridization to examine *Osm* expression in the CNS. SC from EAE mice at 17-dpi showed drastically increased *Osm* mRNA expression, observed particularly in ventral and lateral white matter lesions with corresponding DAPI-staining, reflecting peripheral cell infiltration (Fig. 5c, d). *Osm* mRNA detection was combined with CD11b staining, which indicated that CD11b^+^ cells are a substantial, but not exclusive, source of *Osm* mRNA in white matter lesions (Supplementary Fig. 5f). This approach also revealed that *Clec7a*^*-/-*^ mice with 17-dpi EAE had reduced *Osm* mRNA expression in CD11b^+^ cells (Fig. 5e, f). Because no statistically significant difference in *Osm* mRNA levels in total SC homogenates between WT and *Clec7a*^-/-^ mice was observed (Supplementary Fig. 5g), Dectin-1-mediated *Osm* expression in the SC is most likely specific to CNS-infiltrated myeloid cells in the microenvironment of EAE white matter lesions.

### Osm upregulation by Dectin-1 is Card9-independent

Next, we investigated how *Osm* expression is enhanced by Dectin-1. Card9 is a major signaling protein downstream of Dectin-1^3, 4^ with pathogenic function in EAE (Fig. 2). Interestingly, curdlan upregulates *Osm* expression in *Card9*^*-/-*^, but not *Clec7a*^*-/-*^ myeloid cells (Fig. 6a, Supplementary Fig. 6a), indicating Card9-independent *Osm* expression by Dectin-1. This was unexpected, as the majority of studies of Dectin-1 signaling have focused on Card9-mediated effects. Given the divergent functions for Card9 and Dectin-1 in EAE development (Fig. 1, 2), we hypothesize that Card9-independent signaling may activate a Dectin-1-mediated transcriptional program with distinct protective functions, as seen in driving *Osm* expression.

**Figure 6.**
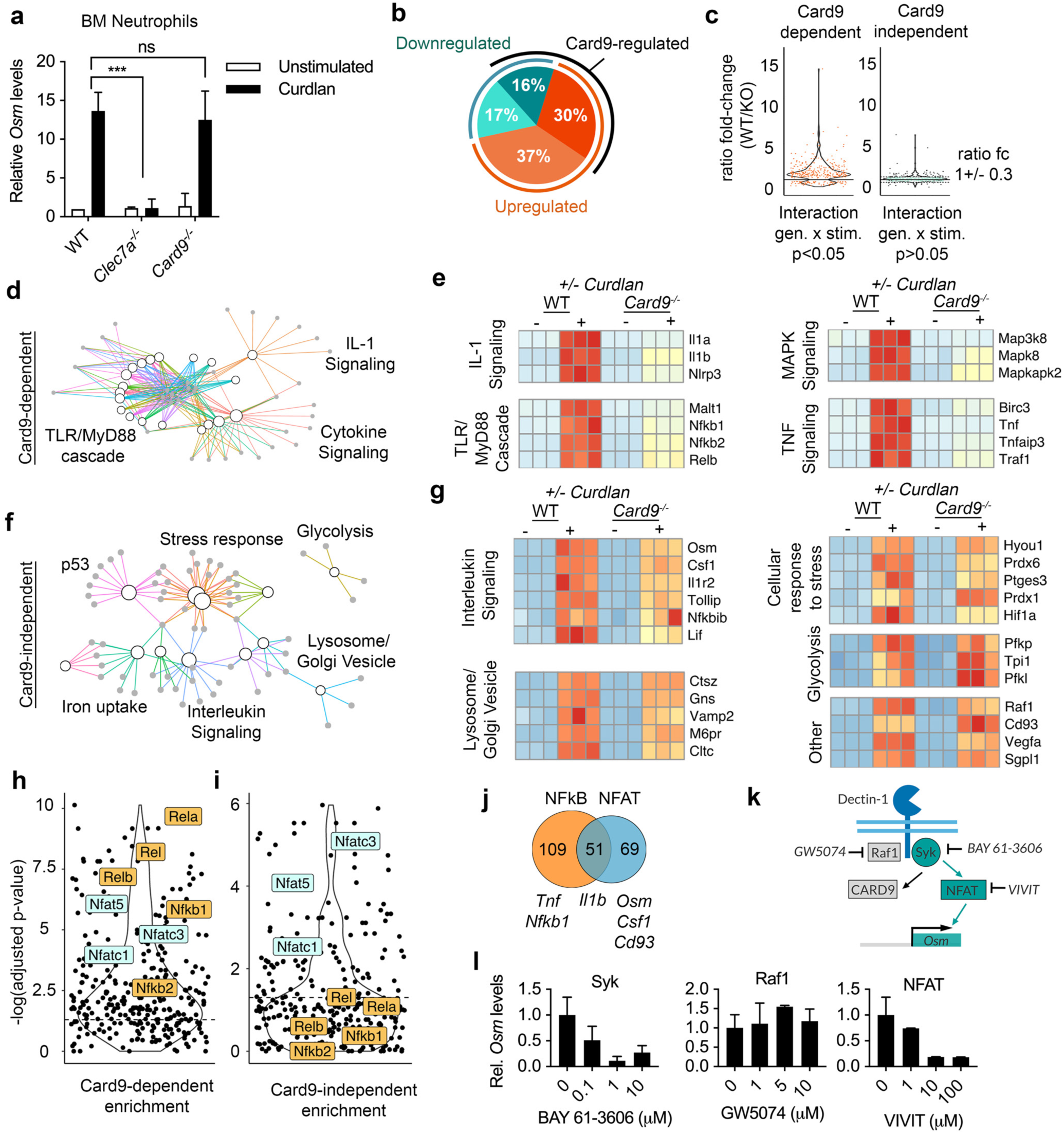
Defining the Card9-independent Dectin-1 transcriptional program. **(a)** *Osm* mRNA levels in BM neutrophils from WT, *Clec7a*^*-/-*^, and *Card9*^*-/-*^ mice. Neutrophils were treated with or without curdlan (100 μg/ml) *ex vivo* for 3 hrs. Mean ± SEM, *n*=3 mice/group. Representative of 3 independent experiments. **(b-g)** RNAseq analyses of BM neutrophils from WT and *Card9*^*-/-*^ mice (*n*=3 mice/group) treated with or without curdlan (100 μg/ml) *ex vivo* for 3 hrs. Proportions of genes, significantly up-or down-regulated in a Card9-dependent manner, in WT neutrophils with curdlan stimulation (b). Ratio of gene expression fold-change, comparing WT and *Card9*^*-/-*^ (KO) neutrophils, with curdlan stimulation (points indicate individual genes) (c). Gene-concept network based on RNAseq results, showing pathway enrichment analysis (d, f). Heatmaps of selected Card9-dependent genes with indicated associated pathways (e, g). Card9-dependent and -independent candidate genes are indicated in (d, e) and (f, g), respectively. **(h, i)** Plots of adjusted p-values for TF binding site enrichment near Card9-dependent genes (h) or Card9-independent genes (i). NFAT family and NFκB family TFs are indicated in turquoise and yellow, respectively. **(j)** Venn diagram of genes with at least 3 predicted NFκB or NFAT binding sites in OCRs within 100kb. **(k)** Schematic of Dectin-1 signaling with small molecule inhibitors. **(l)** RT-qPCR evaluation of *Osm* mRNA levels in WT BM neutrophils pre-treated with inhibitors at the indicated doses for 1 hr before curdlan stimulation (100 μg/ml) for 3 hrs. Data representative of 2 independent experiments.

### Card9-independent signaling promotes non-canonical Dectin-1 effector functions

To further characterize the Card9-independent Dectin-1-mediated transcriptional program, we focused on neutrophils, which express Dectin-1 and are among the most abundant myeloid cells in the spleen (Fig. 4b) and in the CNS during EAE^40^ and can limit late-stage disease^41^. We performed RNA sequencing of BM neutrophils from WT and *Card9*^*-/-*^ mice treated with or without curdlan for 3-hrs. We first identified 1,157 genes which were significantly upregulated or downregulated by curdlan stimulation in WT neutrophils (Fig. 6b). Remarkably, approximately half were not significantly regulated by Card9 (Fig. 6b). Indeed, after further selecting those genes with similar expression between WT and *Card9*^*-/-*^ cells (±30 %), we identified a total of 567 Card9-independent candidate genes (Fig. 6c).

Next, we performed pathway analyses^42^ on genes that are upregulated with curdlan stimulation. Card9-dependent genes showed enrichment in pathways such as TLR signaling, MyD88 signaling, and IL-1 signaling (*Il1b, Nlrp3, Nfkb1, Tnf*) (Fig. 6d, e). Thus, as we anticipated, Card9-dependent genes were enriched for pro-inflammatory factors^3, 4^. In contrast, Card9-independent genes were enriched in pathways including iron uptake and processing, lysosome/Golgi vesicle biogenesis, glycolysis, cellular response to stress, and interleukin signaling (Fig. 6f, g). *Osm* was among the Card9-independent genes found in the interleukin signaling pathway, which includes both neuroprotective factors and negative regulators of pro-inflammatory signaling, such as an inhibitor of TLR signaling (*Tollip*), an inhibitor of NFκB signaling (*Nfkbib*), and the decoy receptor for IL-1 (*Il1r2*) (Fig. 6g). Other Card9-independent genes besides *Osm* with reported neuroprotective functions in EAE, include *Csf1*^43^, *Cd93*^*44*^, and *Vegfa*^45^ (Fig. 6g). We confirmed that these Card9-independent genes are upregulated by curdlan in a Dectin-1 dependent manner (Supplementary Fig. 6b). While our initial RNA-seq analysis was performed using neutrophils, we also observed Dectin-1 mediated upregulation of Card9-independent genes in other myeloid cells including monocytes, macrophages, and DCs with some differences depending on cell-type (Supplementary Fig. 6c, d). In summary, Card9-independent Dectin-1 signaling drives expression of *Osm* and other genes encoding neuroprotective molecules in multiple myeloid cell types.

### NFAT regulates Card9-independent Dectin-1/Osm pathway

Next, we looked for enrichment of predicted transcription factor binding sites near our genes of interest by focusing on open chromatin regions (OCRs) in BM neutrophils using publicly available ATAC-seq data from the Immunological Genome (ImmGen) Project ^46, 47^. We found that Card9-dependent genes had significantly enriched binding sites for NFκB-family transcription factors, including Rel-A, Rel, Rel-B, and NFκB1 (Fig. 6h). Although binding sites for NFAT-family transcription factors were enriched near some Card9-dependent genes, Card9-independent genes showed significant enrichment for NFAT binding sites (Fig. 6h-j). Specifically, NFATc3, NFAT5, and NFATc1 were among the top transcription factors with enriched binding sites near Card9-independent genes in our analysis (Fig. 6i). In the *Osm* locus, multiple predicted NFAT binding sites were identified in OCRs found in both neutrophils and monocytes (Supplementary Fig. 6e). Previous studies have shown that Dectin-1 can indeed induce NFAT activity through Syk/PLCγ/Ca^2+^ signaling^48, 49^, but the transcriptional program downstream of this pathway remains less characterized compared to Card9 signaling.

Next, we used small molecule inhibitors to target signaling pathways downstream of Dectin-1 (Fig. 6k). Although inhibition of Raf1 (GW5074) did not block *Osm* upregulation by curdlan stimulation in neutrophils, inhibitors of Syk (BAY 61-3606) and NFAT (VIVIT) did (Fig. 6l). Both inhibitors showed dose-dependent effects (Fig. 6l), and neither resulted in detectable cellular toxicity, suggesting that Dectin-1 enhances *Osm* expression via Syk and NFAT. Other genes, such as *Csf1* and *Vegfa*, behaved as *Osm* did, but some Card9-independent genes, such as *Cd93*, do not require NFAT for upregulation (Supplementary Fig. 6f). Furthermore, interfering with NFAT signaling by inhibiting calcineurin (CaN) and Phospholipase C (PLC) blocked *Osm* upregulation (Supplementary Fig. 6g, h), indicating that CaN and PLC are involved. Reciprocally, ionomycin upregulated *Osm* expression even in the absence of curdlan stimulation (Supplementary Fig. 6i), suggesting increased intracellular calcium levels are sufficient for *Osm* upregulation. These findings demonstrate that Dectin-1 signaling induces *Osm* expression through a non-canonical Card9-independent pathway, which includes Syk, PLC, Ca^2+^, CaN, and NFAT.

### OsmR expressed on astrocyte limits EAE severity

Previous studies suggested that Osm may be protective in different forms of CNS injury acting through its receptor, OsmR, which forms a heterodimer with gp130^36^. Intraperitoneal (*i*.*p*.) treatment with recombinant Osm was shown to block EAE development^18^ and Osm limited demyelination and promoted remyelination in the cuprizone model^17, 36^. However, the protective mechanism of OsmR signaling in CNS autoimmunity remains largely unknown. Here, we sought to determine which cell types express OsmR in the SC from EAE mice. First, we found that OsmR was expressed on GFAP^+^ astrocyte projections in SC white matter in both naïve and EAE mice (Fig. 7a), as well as on axons (Fig. 7b). OsmR expression in astrocytes was confirmed using *Aldh1l1-eGFP* reporter mice (Supplementary Fig. 7a). In contrast, OsmR was not detected in oligodendrocytes expressing MBP (Fig 7c). We also did not observe significant expression of OsmR on CD45^+^ immune cells (Fig. 7d) or Tmem119^+^ microglia (Fig. 7e), consistent with expression data from ImmGen indicating very low OsmR expression on immune cells (Supplementary Fig. 7c)^46^.

**Figure 7.**
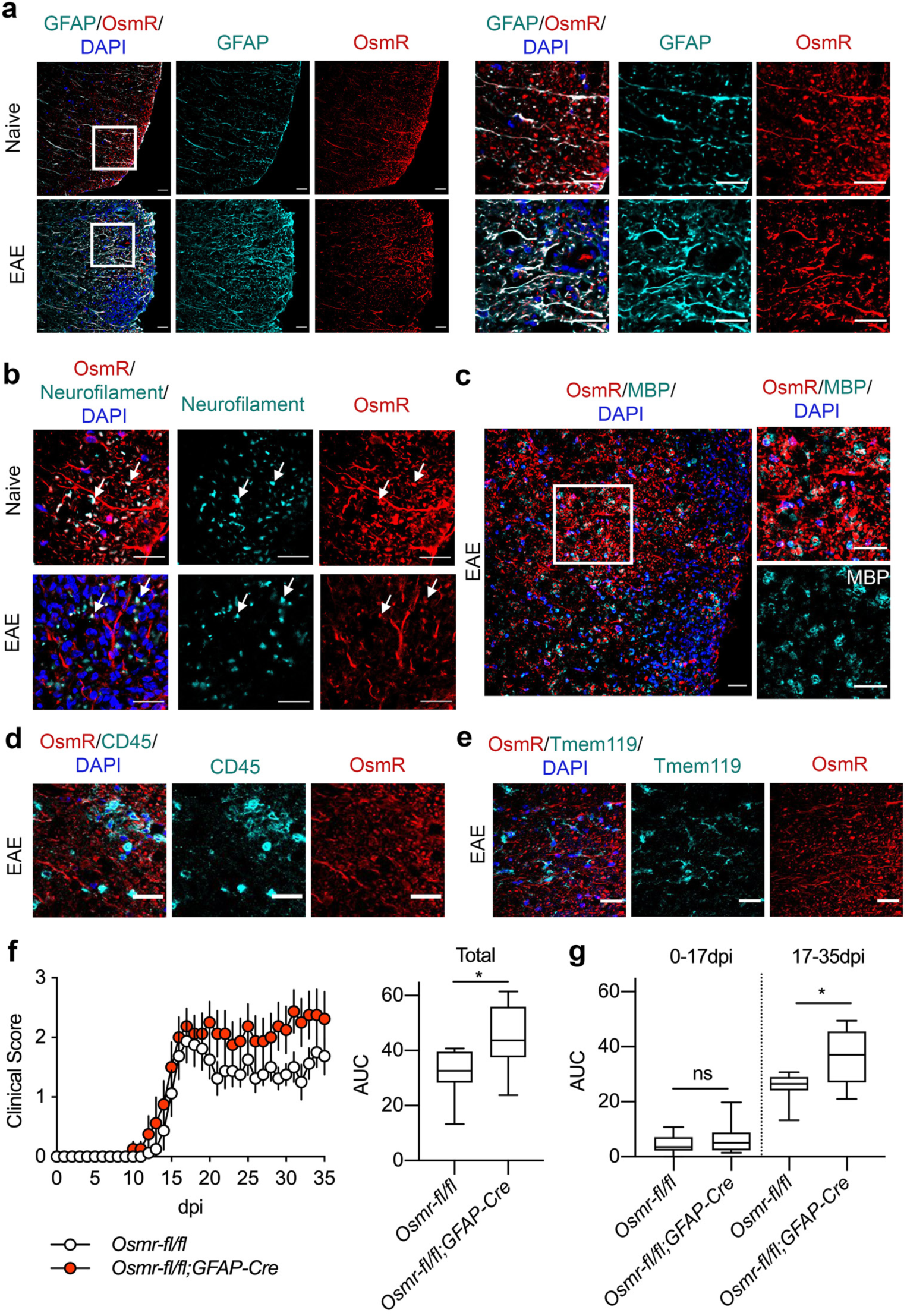
OsmR expression in astrocytes limits EAE severity. **(a-e)** Representative images of OsmR expression in ventro-lateral white matter of lumbar SC from naïve and EAE 17-dpi mice along with counter-staining for GFAP (a), Neurofilament (b), MBP (c), CD45 (d), or Tmem119 (e). Scale, 25μm. Data representative of *n*=5 mice/group combined from two independent experiments. **(f, g)** EAE scores of *Osmr*^*fl/fl*^ and *Osmr*^*fl/fl*^;*Gfap*^*Cre*^ littermates (*n*=8). Data quantified by total AUC (f), or AUC divided into pre-(0-17dpi) and post-peak (17-35dpi) (g). Representative of 3 independent experiments.

Astrocytes can be protective in EAE^50^ and astrocyte expression of gp130, which forms a heterodimer with OsmR, was reported to inhibit EAE pathology^51^. Thus, we focused our investigation on OsmR function in astrocytes using a conditionally deletion approach (Supplementary Fig. 7d-f). We backcrossed *Osmr*^*fl/fl*^ mice^52^ (from The Jackson Laboratory) to C57BL/6J (B6) WT mice using a speed congenic approach and obtained mice with 99.78% of the B6 genomic background (Supplementary Fig. 7g) prior to crossing to *Gfap*^*Cre* 53^. We found that deletion of *Osmr* in astrocytes (*Gfap*^*Cre*^;*Osmr*^*fl/fl*^) exacerbated disease severity compared to littermate controls (*Osmr*^*fl/fl*^) (Fig. 7f). Notably, *Gfap*^*Cre*^;*Osmr*^*fl/fl*^ mice showed impaired remission (day 18-35), although no difference in EAE severity was observed before the disease peak (day 0-17) (Fig. 7f, g). These results indicate that OsmR expressed on astrocytes is protective specifically in the remission phase of EAE. Additionally, we observed Dectin-1^+^ cells and Ly6G^+^ neutrophils in close proximity to GFAP^+^ astrocytes in the SC of EAE mice (Supplementary Fig. 7h, i), suggesting the potential for crosstalk between myeloid cells and astrocytes. In summary, we found that non-canonical Dectin-1 signaling promotes expression of neuroprotective and anti-inflammatory factors in myeloid cells including *Osm*, and OsmR in astrocytes is protective in CNS autoimmunity.

## DISCUSSION

In this study, we found that the CLR Dectin-1 limits CNS inflammation in a mouse model of MS through a non-canonical protective pathway. Dectin-1 is generally known for its pro-inflammatory response in the setting of fungal infection, particularly by inducing IL-1β via Card9/NFκB signaling and subsequent Th17 responses^3, 4, 5^, which are considered to be highly encephalitogenic^24^. Thus, the protective role of Dectin-1 in EAE was initially unexpected. We found that non-canonical Card9-independent Dectin-1 signaling induces expression of the neuroprotective cytokine, *Osm*, and a transcriptional program with protective and anti-inflammatory functions. In humans, eQTLs regulating *CLEC7A* (Dectin-1) gene expression have been identified^54, 55^ and *CLEC7A* is expressed in active MS lesions (Supplementary Fig. 1)^20, 21^. Despite the well-known inflammatory role of Dectin-1, some studies have suggested tolerogenic and anti-inflammatory functions of Dectin-1 signaling^9, 56, 57^. Furthermore, Dectin-1 signaling promotes axon regeneration following optic nerve crush injury^34^ and treatment with zymosan (TLR2/Dectin-1 agonist) reduced EAE severity^58^. Thus, harnessing protective non-canonical Dectin-1 signaling merits further investigation as a potential therapeutic approach for MS.

Studies on CLRs in EAE and related models have mostly focused on inflammatory responses to hk*Mtb* as an adjuvant, ligands corresponding to CLRs of interest^13, 28, 59, 60^, and endogenous ligands^61^. We demonstrate that Dectin-1 does not detect hk*Mtb*. Furthermore, protective Dectin-1 functions in EAE are unlikely to be attributable to recognition of commensal fungi, because Dectin-1 limited EAE severity even with administration of an oral antifungal agent (*data not shown*). While we cannot completely rule out the contribution of microbial ligands, our findings suggest involvement of endogenous Dectin-1 ligands, such as Galectin-9^6^, vimentin^7^, Annexin proteins^9^, and various unidentified proteins with N-glycans^29^. We are currently studying the impact of these endogenous Dectin-1 ligands on EAE.

We found that the protective function of Dectin-1 was mediated by hematopoietic-derived cells. Yet, microglia also upregulated Dectin-1 expression in EAE, consistent with the disease-associated microglia (DAM) phenotype^62^. Given the limitations of the BM chimera approach, we cannot fully rule out a contribution of microglial Dectin-1 in EAE. Nevertheless, hematopoietic-derived myeloid cells including neutrophils, monocytes, and DCs are likely to be important mediators of Dectin-1 function in EAE. Neutrophils in particular may be involved, given their abundance^40^, protective functions^41^, and localization at sites of demyelination and axonal damage in the SC parenchyma during EAE63, 64, 65 66, 67. While the role of neutrophils in MS remains elusive, multiple studies suggest involvement of neutrophils, or granulocytic myeloid-derived suppressor cells (G-MDSCs), in both MS and EAE^41, 68, 69, 70, 71, 72, 73, 74, 75, 76^. Here, further studies using a conditional deletion approach will be essential to determine which myeloid cells mediate protective functions of Dectin-1 in EAE.

We found that Dectin-1 promotes expression of Osm, a cytokine with described protective functions in cuprizone-induced demyelination^17, 77^ and other models of neuropathology^16, 37^. Osm protein is expressed in MS brains^38^ and is elevated in supernatants of cultured PBMCs from MS patients^39^. In addition, a SNP in the *OSMR* gene was identified as a candidate risk factor for MS^78^. In this study, we found that *Osm* is expressed by myeloid cells in the SC of EAE mice and is upregulated by Dectin-1 signaling. Furthermore, we found that OsmR in astrocytes is protective in the remission phase of EAE. Astrocytes are increasingly recognized as important players in CNS autoimmunity^79, 80^. In particular, understanding the mechanisms of crosstalk between myeloid cells and astrocytes through the Osm-OsmR axis could address a significant unmet need in MS: halting CNS injury and promoting recovery.

This study contributes to a growing body of evidence that Dectin-1 has important roles beyond the context of fungal infection. We advance this emerging understanding by identifying a protective mechanism of Dectin-1 specifically in the setting of CNS autoimmunity. Together, our findings provoke a reconsideration of Dectin-1 and its functions beyond the context of infection and provide potential novel targets for therapeutic intervention in neuroinflammatory disorders.

## MATERIALS AND METHODS

### Mice

Both male and female mice of the C57BL/6 (B6) background were used in this study. Dectin-1 deficient (*Clec7a*^*-/-*^) mice were originally generated by Dr. Gordon Brown (University of Aberdeen)^5^. *Card9*^*-/-*^ mice were provided by Dr. Xin Lin (M.D. Anderson Cancer Center, Houston, TX) ^81^. *Osmr-fl* mice on a mixed C57BL/6 and 129×1/SvJ background (JAX #011081) were rederived from a cryopreserved line at Jackson Laboratories^52^. The N1 generation following the first backcross to B6 in our lab showed 92.4-95.4 % of B6 genomic background, and further backcrossed to B6 for 5 additional generations using marker-assisted speed congenic approach (DartMouse) until we achieved 99.8 % of the B6 genomic background (Supplementary Fig. 7g). *Osmr-fl* mice were then crossed to *Gfap*^*Cre*^ (JAX #024098) ^53^, and littermate controls were used for *Osmr-fl* experiments. Fixed tissues from *Aldh1l1-eGFP* mice (MMRRC #001015-UCD) (originally generated by the laboratory of Dr. Ben Barres at Stanford University)^82^ were provided by the laboratory of Dr. Cagla Eroglu (Duke University). Six-to eight-week old age- and sex-matched mice were used for experiments unless otherwise specified. All mice were housed under specific pathogen-free conditions and all animal experiments were performed as approved by Institutional Animal Care and Use Committee at Duke University.

### Reagents, antibodies, and recombinant proteins

Curdlan (β-1,3 glucan hydrate, from *Alcaligenes faecalis*, Sigma-Aldrich) was used as a Dectin-1 specific agonist for *ex vivo* experiments, unless otherwise indicated. Hot-alkali treated depleted zymosan (dz) (InvivoGen) was used as a Dectin-1 specific ligand for *in vivo* experiments. Heat-killed *Mycobacterium tuberculosis* (hk*Mtb*) H37a Ra was purchased from BD Difco™. Small molecule signaling inhibitors were purchased from Cayman Chemical, with the exception of GW5074 (Sigma Aldrich), and concentrations for their use were determined using supplier recommendations and based on previous publications. Recombinant mouse (rm) M-CSF and GM-CSF proteins were obtained from BioLegend. MOG_35-55_ peptide (MEVGWYRSPFSRVVHLYRNGK) was synthesized by United Biosystems. Enzyme-linked immunosorbent assay (ELISA) kits for detection of mouse Osm were purchased from R&D. Detailed list of reagents can be found in Supplementary Table 1.

### EAE induction and evaluation

Unless otherwise noted, EAE was induced as follows: MOG_35-55_ peptide (100 μg in 100 μl PBS) emulsified in CFA (100 μl including 2 mg/ml hk*Mtb*) was subcutaneously (*s*.*c*.) injected into lower flanks of mice. Intraperitoneal injection of pertussis toxin (List Biologicals; 200 ng in 200 μl PBS) was performed on day 0 and 2. EAE induction was performed with reduced adjuvant in Fig. 1b and Fig. 4f, g by using 0.5 mg/ml instead of 2 mg/ml hk*Mtb* CFA (50 μg/mouse instead of 200 μg/mouse). Mice were monitored daily for clinical signs of EAE and scores were assigned based on the following criteria: 0.5, partial tail limpness; 1, tail limpness; 1.5, impaired righting reflex; 2, partial hindlimb paralysis; 2.5, partial hindlimb paralysis with dragging of at least one hind paw; 3, bilateral hindlimb paralysis; 3.5, severe bilateral hindlimb paralysis with hunched posture; 4, hind- and forelimb paralysis; 5, death.

### Statistical analysis

EAE clinical scores were analyzed by calculating the area under the curve (AUC) summing clinical scores over the indicated period. Statistical analysis of EAE clinical score AUC was performed using a non-parametric Mann-Whitney U-test and, where indicated, AUC was normalized to the mean of the control group. Box plots of EAE clinical score AUC data include the following elements: center line, median; box limits, upper and lower quartiles; whiskers, minimum and maximum values. For remaining analyses, student’s t-test (paired or unpaired, as appropriate, two-sided in all cases) or analysis of variance (ANOVA) was applied (one- or two-factors, repeated measures as appropriate) as indicated in figure legends. If a two-factor ANOVA yielded an interaction term with *p<0*.*1*, then post-hoc Sidak testing with correction for multiple comparison was applied. If a two-factor ANOVA did not result in a significant interaction term (*p>0*.*1*), then any significant main effects of interest are indicated in the figure and legend. Log-transformation was performed prior to 2-factor ANOVA as appropriate when analyses of flow cytometry data included cell numbers that varied greatly in magnitude between multiple cell types (Fig. 2d, e; Supplementary Fig. 2c, d; Supplementary Fig. 3a, b). All results are expressed as means ± SEM unless otherwise noted, and criterion of significance was set as *p<0*.*05*(*), *p<0*.*01*(**), or *p<0*.*001*(***).

### Cell isolation for flow cytometry analysis and FACS-sorting

Mice were euthanized with CO_2_ in addition to a secondary method. BM was isolated from the femur and tibia by flushing marrow with a needle and syringe filled with sterile PBS and pipetting to obtain a single cell suspension. Spleens and ILNs were dissected and homogenized using sterile glass slides. SCs were isolated by flushing the SC from the vertebral column with a needle and syringe filled with sterile PBS, minced in PBS including 5% FBS and 1mM HEPES, then digested with Collagenase D (Roche) at 37 C for 30 minutes. Following digestion, single cell suspensions were prepared from SC tissues by passing through an 18G needle and filtered through a 70 μm cell strainer. Cells were then resuspended in 38% isotonic Percoll and centrifuged at 2,000 g for 30 min with no brake. Following centrifugation, the lipid and debris layer was aspirated from the top of the tube and the cell pellet was resuspended. All tissues were treated with RBC lysis solution prior to staining for flow cytometry.

### Fluorochrome-conjugated antibody staining for flow cytometry

Staining for live cells was performed using a LIVE/DEAD™ fixable dead cell stain kit (Invitrogen) prior to staining with fluorochrome-conjugated antibodies listed in Supplementary Table 3. Cells were analyzed by flow cytometry using either Fortessa X20 or FACSCanto (BD systems). Flow cytometry results were analyzed using FlowJo. Gating for specific cell types was performed using the following markers after pre-gating on live cells: neutrophils (CD11b^+^Ly6G^+^), monocytes (CD11b^+^Ly6G^-^Ly6C^hi^), cDCs (CD11b^+^CD11c^+^), F4/80^hi^ macrophages (CD11b^+^F4/80^hi^), B cells (CD19^+^), CD4^+^ T cells (CD3^+^CD4^+^), CD8^+^ T cells (CD3^+^CD8^+^), microglia (CD45^lo^CD11b^lo^ and Tmem119^+^, as indicated in the text).

### Intracellular cytokine staining for flow cytometry

Intracellular cytokine staining was performed on spleen and LN single cell suspensions following 5-6 hr stimulation with PMA (10ng/ml) (Sigma Aldrich) and ionomycin (1μg/ml) (Sigma Aldrich), with GolgiPlug (BD), added for the final 2 hrs of stimulation. Staining performed using a LIVE/DEAD™ fixable dead cell stain kit (Invitrogen) followed by fluorochrome-conjugated antibodies for cell surface proteins before fixation and permeabilization (BD Cytofix/Cytoperm Kit). Antibody staining for intracellular cytokines was then performed in permeabilization buffer. Intracellular staining for Foxp3 was performed on single cell suspensions from spleen and LNs specifically using a FOXP3 Fix/Perm Kit (BioLegend).

### Isolation of neutrophils and monocytes and BM-derived cell culture

BM neutrophils were isolated using MACS-column purification (Millentyi Biotec) with biotinylated mouse Ly6G antibodies (BioLegend) and streptavidin magnetic beads (Millentyi Biotec) for *ex vivo* study. For comparisons of BM neutrophils and BM monocytes, cells were isolated by FACS-sorting on CD11b^+^Ly6G^+^ (neutrophils) and CD11b^+^Ly6G^-^Ly6C^+^ (monocytes) cells (Astrios sorter, Beckman Coulter). BMDMs were generated by culturing BM cells with rmM-CSF (20 ng/ml, BioLegend) and BMDCs were generated with rmGM-CSF (20 ng/ml, BioLegend). We used complete RPMI medium for all cell culture studies.

### Isolation of spinal cord tissue for histology

Transcardial perfusion was used to maximize integrity of brain and spinal cord tissue to improve sample quality. Specifically, mice were administered sodium pentobarbital (100 mg/kg) by intraperitoneal injection and monitored until complete anesthesia was achieved (non-responsive to pedal reflex). The thoracic cavity was dissected, and the inferior vena cava severed prior to insertion of a needle in the left ventricle and slow manual administration of PBS followed by 4% paraformaldehyde (PFA). Following transcardial perfusion and fixation, spinal cord tissue was manually dissected and fixed in 4% PFA overnight before cryoprotection in 30% sucrose for 1-2 days. Tissues were embedded in Tissue-Tek OCT compound (Sakura), frozen, and stored at -80 °C.

### LFB-PAS staining and analysis

Luxol fast blue (LFB)-Periodic Acid Schiff (PAS) staining was performed on transverse fixed-frozen sections (10 μm) of lumbar spinal cord using LFB Solvent blue 38, Gill’s Hematoxylin No. 3, and Schiff’s reagent (all obtained from Sigma-Aldrich). LFB-PAS stained sections were imaged on the Zeiss Axio Imager widefield microscope.

### Immunofluorescent (IF) staining and imaging

Staining was performed with either floating sections in tissue culture wells or thaw-mounted sections on slides. Transverse fixed-frozen sections (25 μm) of lumbar spinal cord were washed with TBS-T, permeabilized with TBS 0.25% Triton 1Î for 15min, washed with TBS-T, then incubated in blocking buffer (TBS-T 3% BSA) for at least 1 hr. Primary antibody staining was performed overnight at 4 °C, followed by washing in TBS-T. Secondary antibody staining with fluorochrome-conjugated antibodies was performed for 2 hrs at 4 °C (antibodies listed in Supplementary Table 3). Stained sections were washed and mounted using ProLong Gold Antifade Mountant with DAPI (ThermoFisher). Slides were imaged using the Zeiss 710 inverted confocal microscope and analyzed using ImageJ. IF images are typically representative of at least two hpf/sample and data was obtained from several individual animals per experiment (sample size (*n*) denoting number of mice).

### Dectin-1 reporter assay

To generate a Dectin-1 reporter system, we used a chimeric mouse Dectin-1 receptor (extracellular and transmembrane domains) fused to the cytoplasmic tail of mouse CD3ζ, cloned into the retroviral pMXs-IP-mCD3ζ-mDectin1 vector^26^, a gift from Gordon Brown (University of Aberdeen, UK) under MTA). pMXs-IP-mCD3ζ-Dectin-1, together with Pcl-Eco (packaging vector), was transfected to BOSC cells using lipofectamine LTX. The 58α^-^β^-^ mouse T hybridoma cell line with stable expression of a NFAT-GFP-hCD4 RV reporter construct^27^ (gift from Ken Murphy, Washington University) was retrovirally transfected to express the Dectin-1 chimeric protein. Activated Dectin-1 was reported by GFP expression by the hybridoma and detected by flow cytometry.

### mRNA detection in tissue sections by RNAscope

To prepare samples for RNAscope, lumbar spinal cord (SC) segments were dissected following transcardial perfusion (described above) and post-fixed overnight in 4% PFA. Samples were then transferred to 30% sucrose in PBS and kept overnight. Spinal cords were cryosectioned to 20 μm and thaw-mounted onto Superfrost Plus slides (Fisher Scientific). Slides were allowed to dry for 20 minutes at room temperature, and then stored at -20 C overnight. The RNAscope Multiplex Fluorescent Reagent Kit v2 (Advanced Cell Diagnostics, ACD) was used for *in situ* hybridization as previously described^83^. Tissue pretreatment was performed for 30 min with Protease IV in the RNAscope kit at RT followed by probe hybridization and detection according to manufacturer’s instructions. We used probes designed by Advanced Cell Diagnostics to *Osm* (NM_001013365.2), specifically Mm-Osm (#427071). IF staining with antibody to CD11b was performed after the RNAscope procedure using methods described above. Slides were imaged using the Zeiss 710 inverted confocal microscope. For comparison of naïve and EAE conditions, *Osm* mRNA puncta were quantified by performing object detection using ImageJ, calculating the integrated density for each punctum, and summing over all puncta in a given 350 μm^2^ high-power field (hpf). Positive signal, *i*.*e*., puncta, was defined as >5-fold higher than background. Five hpf per mouse were analyzed, one in each indicated region of lumbar spinal cord. For evaluation of *Osm* expression in CD11b^+^ cells, z-stack images with 1.16 μm spacing were collected through the full thickness of each section. For each 350 μm^2^ ventrolateral white matter hpf per mouse, *Osm* puncta were quantified manually in a blinded manner. Specifically, *Osm* puncta per CD11b^+^ cell were quantified by examining multiple z-stacks in 3D space to identify cell-associated puncta.

### RT-qPCR analysis

To evaluate gene expression in *ex vivo* stimulation experiments, total RNA was extracted from cells with TRIzol on purified cells, while RNeasy Micro Kit (Qiagen) was used for FACS-sorted cells. cDNA synthesis was performed using qScript cDNA Mix (Quantabio). Real-time, quantitative polymerase chain reaction (RT-qPCR) was performed with SYBR FAST qPCR Master Mix (Kapa Biosystems) with primers shown in Supplementary Table 2. Relative amounts of qPCR product were determined using the -*ΔΔCt* method^84^ comparing relative expression of housekeeping (*Actb)* and target genes. Error bars denote mean ± SEM of biological replicates in Fig. 3d; Fig. 5a, b; Fig. 6a; and Supplementary Fig. 5g. In Figures 6l; Supplementary Fig. 5a, d and Supplementary Fig. 6a-d, f error bars (sometimes too short to be identified) were calculated using RQ-Min = 2^-(*ΔΔCt +* T *SD(*ΔCt*))^ and RQ-Max = 2^-(*ΔΔCt -* T *SD(*ΔCt*))^ from triplicate wells as suggested by Applied Biosystems (manufacturer of qPCR machines). Results shown are representative of multiple independent experiments with similar results.

### RNA-seq and analysis

BM neutrophils were obtained by MACS-column purification as described above. Each sample was obtained from an individual WT or *Card9*^*-/-*^ mouse (*n*=3). Cells were stimulated for 3 hrs with curdlan (100 μg/ml). Total RNA was extracted from cells using the RNeasy Micro Kit (Qiagen). mRNA libraries were prepared using the KAPA stranded mRNA-seq kit by Duke Sequencing and Genomic Technologies Shared Resource facility. RNA libraries were sequenced on an Illumina HiSeq4000 using 50-bp single reads.

Initial processing and analysis of sequencing results was performed by Duke Genomic Analysis and Bioinformatics core facility as described here: RNA-seq data was processed using the TrimGalore toolkit which employs Cutadapt^85^ to trim low-quality bases and Illumina sequencing adapters from the 3’ end of the reads. Only reads that were 20nt or longer after trimming were kept for further analysis. Reads were mapped to the GRCm38v73 version of the mouse genome and transcriptome^86^ using the STAR RNA-seq alignment tool^87^. Reads were kept for subsequent analysis if they mapped to a single genomic location. Gene counts were compiled using the HTSeq tool. Only genes that had at least 10 reads in any given library were used in subsequent analysis. Normalization and differential expression was carried out using the DESeq2^88^ Bioconductor^89^ package with the R statistical programming environment^90^ (Supplementary Table 4). The false discovery rate was calculated to control for multiple hypothesis testing.

### Identification of Card9-dependent and -independent candidate genes

We first selected genes with significant differential expression following curdlan stimulation in WT neutrophils (log_2_ fold-change >|1.5| and adjusted p-value<0.05). Among significantly differentially expressed genes, we then selected Card9-dependent candidate genes (adjusted p-value <0.05 for interaction of genotype and stimulation). Among the remaining genes, (adjusted p-value >0.05 for interaction of genotype and stimulation), we further selected those genes with a stimulation-induced fold-change within 30 % of WT values (1 ± 0.3) and defined these as Card9-independent candidate genes (Supplementary Table 5).

### Reactome pathway enrichment analysis

Using these Card9-dependent and -independent candidate gene lists, we performed pathway enrichment analysis on those genes that are upregulated with curdlan stimulation using the *ReactomePA* package and gene set overlap analysis^42^ (Supplementary Table 5) and plotted the resulting enriched pathways (adjusted p-value <0.05) using a *cnetplot* network diagram (genes are small grey points, pathways are colored, lines indicate correspondence of genes with pathways).

### Transcription factor enrichment analysis

To identify potential transcription factors that may regulate Card9-dependent and -independent candidate genes, we developed an analysis approach which leverages existing datasets and tools. Our goal was to identify potential TF candidates based on predicted binding sites near genes in either the Card9-dependent or -independent candidate gene lists. We limited our analysis of predicted binding sites to OCRs in our cell type of interest (BM neutrophils) to better target relevant sites for possible TF binding. We obtained analyzed ATAC-seq result files from Immgen for BM neutrophils which contained information on each OCR, an OCR score for the particular cell type, predicted TF binding sites within each OCR, and genes within 100kb of each OCR^46, 47^ (downloaded from Immgen site on May 22^nd^, 2019). From this genome-wide dataset, we selected only OCRs with a score of >10 in BM neutrophils to specifically include regions of open chromatin in our analysis. From these OCRs, we generated a paired list of TFs (with predicted binding sites within a given OCR) and genes (within 100 kb of a given OCR). This paired list was then used as a custom reference for gene set overlap analysis, in which the category is a given TF. Then we used *GOseq*^91^ to perform gene set overlap analysis between either Card9-dependent or - independent genes and our custom reference. Benjamini Hochberg adjusted p-values were obtained for each TF category for either Card9-dependent or -independent genes (Supplementary Table 5). We then plotted the -log(adjusted p-value) for each TF on a violin plot and labeled TFs of interest in the NFκB or NFAT families (Fig. 6h, i). In addition, we plotted a Venn diagram of genes with at least 3 predicted binding sites of either NFκB or NFAT family TFs (within 100 kb of gene) (Fig. 6j)

## SUPPLEMENTARY MATERIALS

Supplementary Table 1 – Key reagents

Supplementary Table 2 – Primer sequences

Supplementary Table 3 – Antibodies

Supplementary Table 4 – RNA-seq differential expression analysis results

Supplementary Table 5 - Additional analysis of Card9-independent and -dependent candidate genes

## Acknowledgements

Funding: This study was funded by the National Multiple Sclerosis Society (Pilot Grant PP-1509-06274; Research Grant RG 4536B2/1) to M.L.S. and the NIH (F30 AI140497) to M.E.D. M.E.D. is a recipient of the Gertrude B. Elion Mentored Medical Student Research Award of the Triangle Community Foundation.

We appreciate the assistance of the Duke Center for Genomic and Computational Biology core facility with RNA sequencing, alignment, and differential expression analysis. We also appreciate the Duke Cancer Institute Flow Cytometry Core for performing flow cytometry-based sorting. Christabel Tan in the laboratory of Cagla Eroglu, PhD provided samples from *Aldh1l1-eGFP* mice. The laboratory of Gordon Brown (University of Aberdeen, UK) provided the pMXs-IP-mCD3ζ - mDectin1 construct under MTA. The laboratory of Ken Murphy (Washington University) provided the 58α^-^β^-^ mouse T hybridoma cell line with stable expression of a NFAT-GFP-hCD4 RV reporter construct. Megumi Matsuda, MD, Qianru He, MD, and Zilong Wang, PhD in the laboratory of Ru Rong Ji, PhD provided assistance with RNAscope *in situ* hybridization. Tomoko Kadota and Tamira-Marie Bickems provided assistance with mouse genotyping.

## Author contributions

MED, KD, MI, and MLS designed experiments, analyzed and interpreted data. MED performed experiments apart from the following contributions: KD and MI performed experiments in Fig. 1a-b; d-f and Supplementary Fig. 2b-d. KD performed experiments in Fig. 3b-c and Supplementary Fig. 3h. TN assisted with experiments in Fig. 7a-e. EC assisted with experiments in Supplementary Fig. 6b, d. NA assisted with experiments in Fig. 1c. WB provided assistance with EAE methods and experiment design. RRJ provided support with RNAscope *in situ* hybridization. MED and MLS wrote the manuscript with editing by the other co-authors.

## Data and materials availability

Relevant data and materials from this study will be made available upon request. RNA sequencing results will be made available in GEO database under the accession number GSE148850.

## Code availability

All custom code used in this study will be made available upon request through GitHub.

---

**Supplementary Figure 1.**
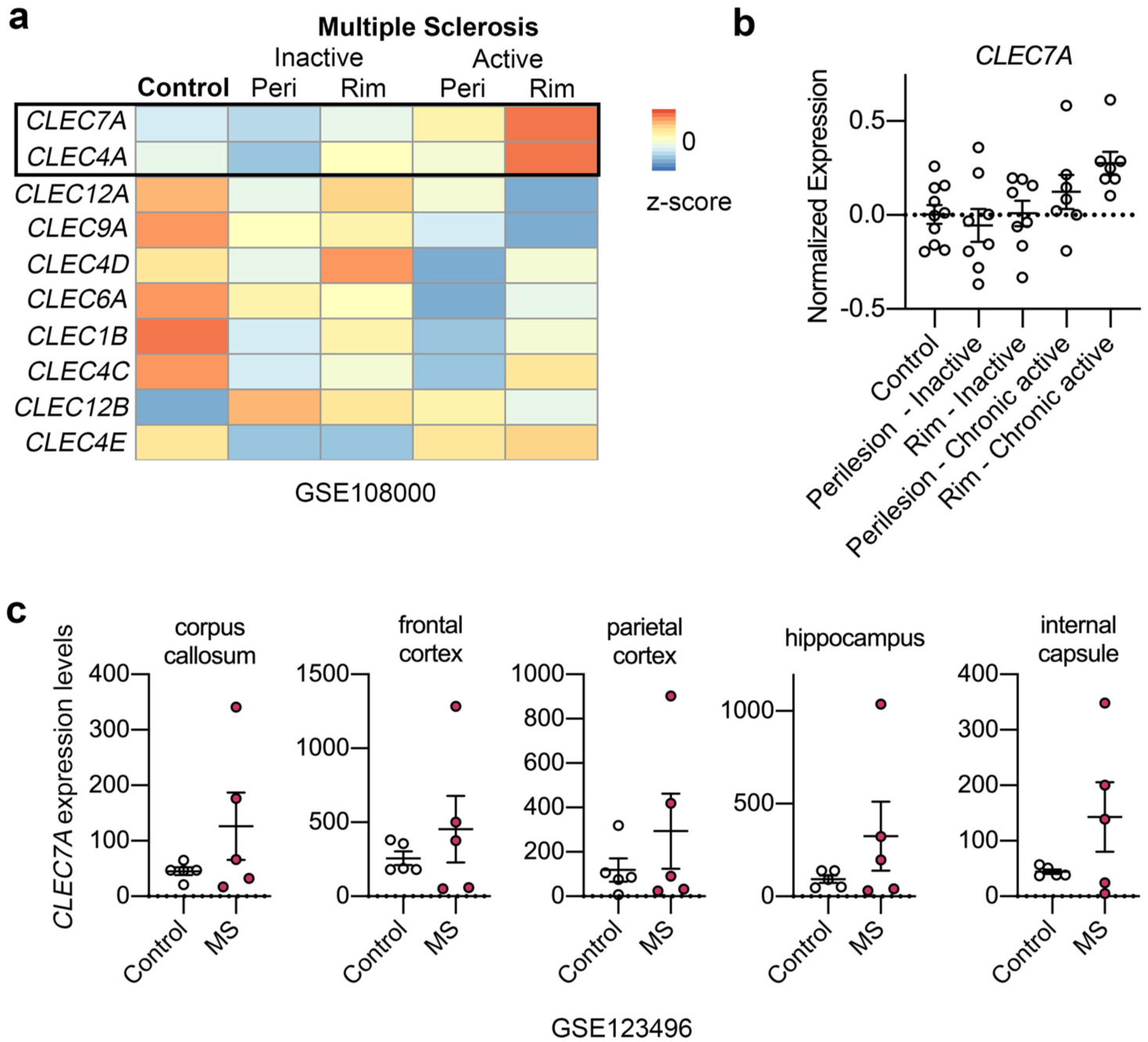
Elevated gene expression of *CLEC7A* in the brains of MS patients. **(a, b)** Gene expression patterns in the rim and peri-lesional normal appearing white matter (NAWM) from MS patients. Heatmap of row-normalized z-scores for expression of selected genes encoding CLRs (a). *CLEC7A* individual normalized expression values (b). Data was obtained by reanalyzing microarray results from GSE108000, including a total of 7 chronic active MS lesions, 8 inactive MS lesions, and WM of 10 control donors. **(c)** *CLEC7A* mRNA levels in indicated brain regions. Data was obtained by reanalyzing RNA-seq results from GSE123496, including a total five MS patients and five age-matched healthy control donors.

**Supplementary Figure 2.**
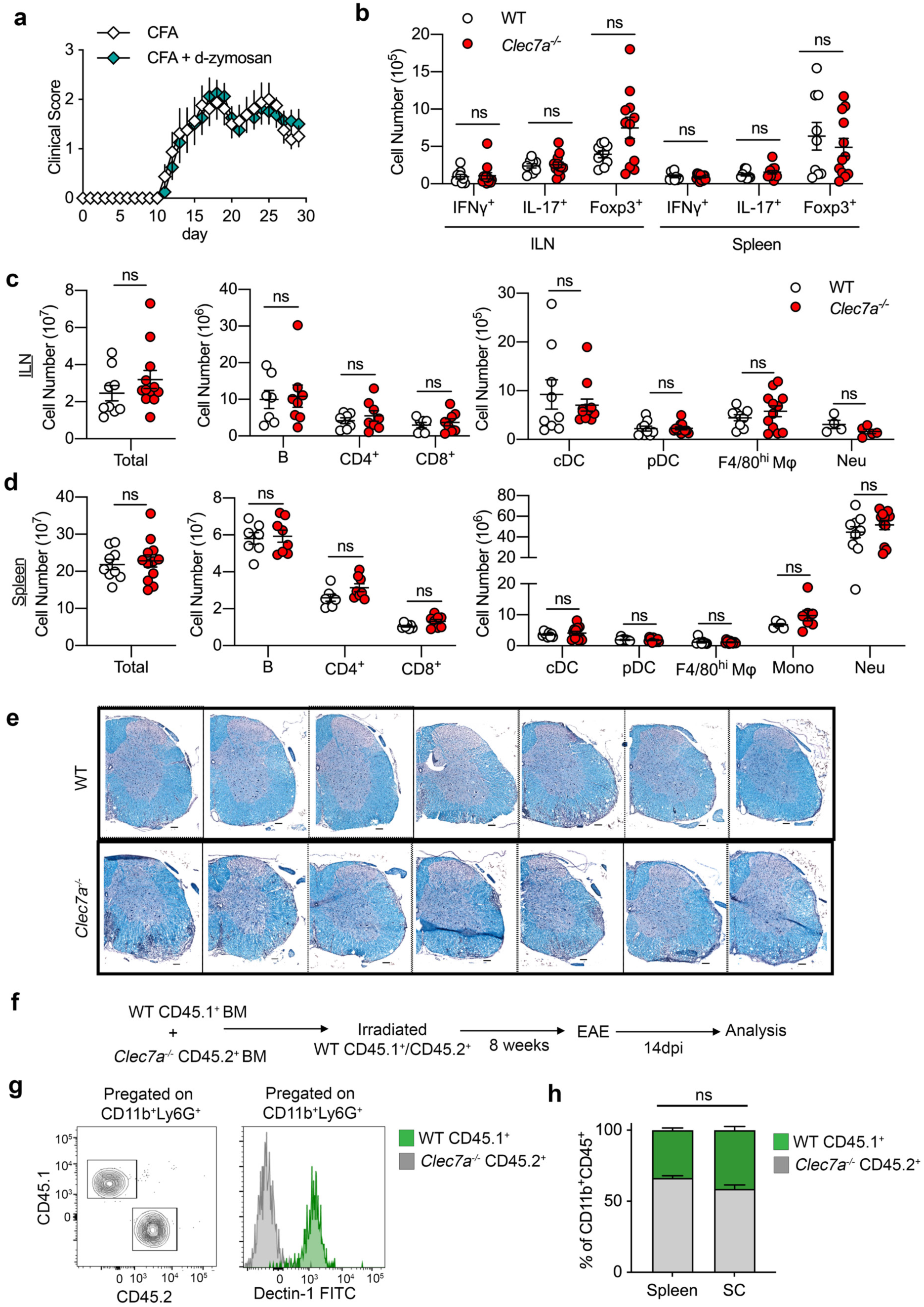
Immunophenotyping of WT, *Clec7a*^*-/-*^, and chimeric mice in EAE. **(a)** EAE clinical scores in WT mice immunized with subcutaneous injection of standard CFA (200μg hk*Mtb*/mouse) or CFA containing d-zymosan (30 μg/mouse) (*n*=8). Data are combined from two independent experiments. **(b-d)** Cell numbers assessed by flow cytometry in WT and *Clec7a*^*-/-*^ mice at 9-dpi in ILN (b, c) and spleen (b, d). Data are combined from two independent experiments. One datapoint denotes a result from one mouse. **(e)** LFB-PAS staining of lumbar SC in WT and *Clec7a*^*-/-*^ mice at EAE 17-dpi. Each image is from an individual mouse. Scale 100μm. **(f)** Schematic of mixed BM chimera experiment: a 50/50 mixture of BM cells from CD45.1 *Clec7a*^*+/+*^ and CD45.2 *Clec7a*^*-/-*^ mice was transferred into irradiated CD45.1/CD45.2 *Clec7a*^*+/+*^ recipients, EAE was induced 8 weeks after BM cell transfer. **(g)** Flow cytometry analysis of chimerism in splenocytes from EAE 14-dpi mixed chimera mice, showing CD45.1 and CD45.2 staining along with Dectin-1 staining, gated on CD11b^+^Ly6G^+^ cells as a representative myeloid cell population. **(h)** Comparison of frequency of WT CD45.1^+^ and *Clec7a*^*-/-*^ CD45.2^+^ cells out of total CD11b^+^CD45^+^ cells between spleen and SC. *n=*5 chimera mice/group. Mean ± SEM shown. Data is representative of 2 independent experiments.

**Supplementary Figure 3.**
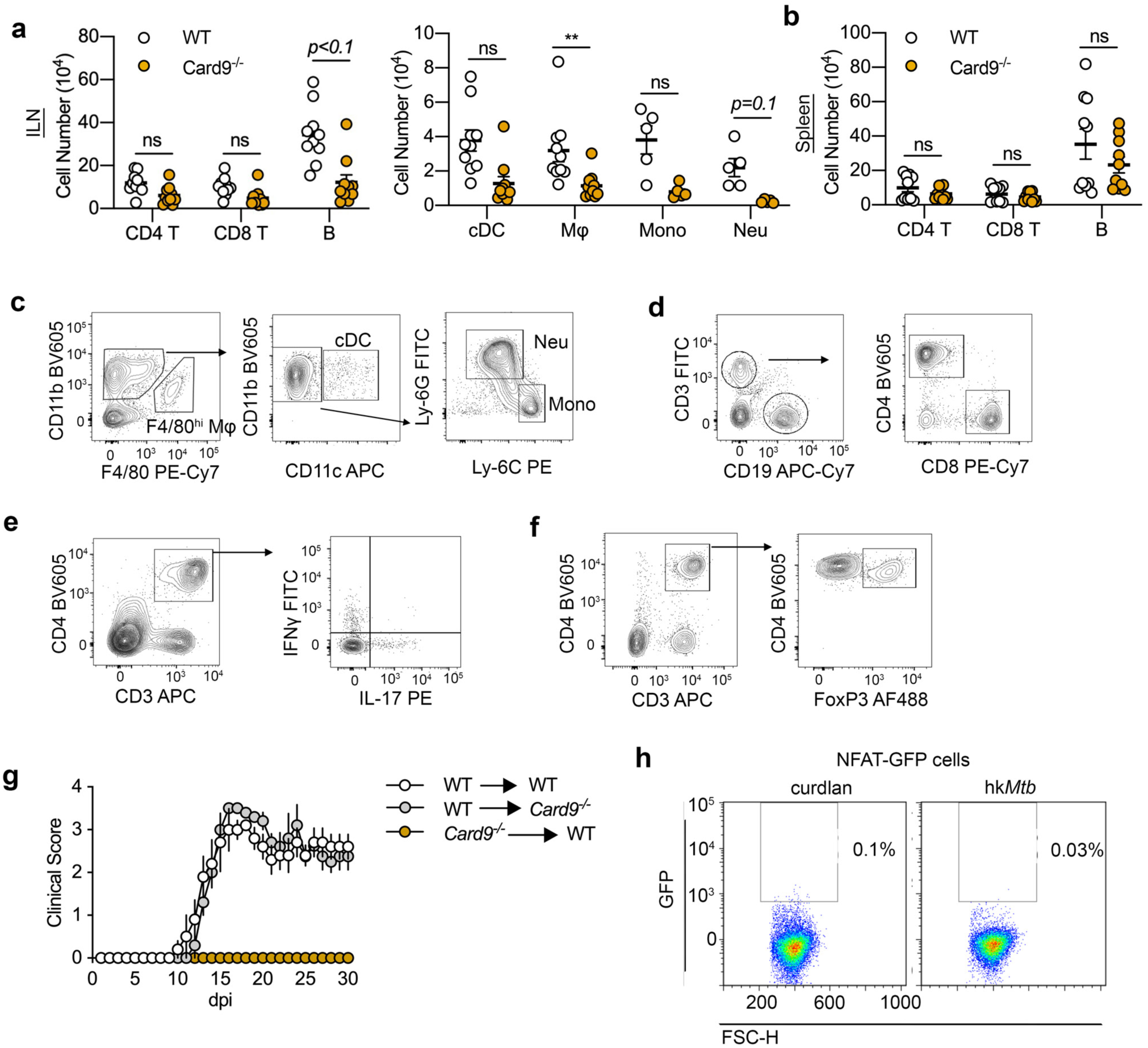
Immunophenotyping of WT and *Card9*^*-/-*^ in EAE and validation of NFAT-GFP reporter cell line. **(a, b)** Cell numbers assessed by flow cytometry in WT and *Card9*^*-/-*^ mice at 9-dpi EAE in ILN (a) and spleen (b). Data are combined from two independent experiments. One datapoint denotes a result from one mouse. **(c-f)** Flow cytometry gating strategy for splenic myeloid cell populations (c), lymphocytes (d), and intracellular cytokine staining for IFNγ, IL-17 (e), and FoxP3 (f). **(g)** EAE clinical scores from BM chimeras. WT BM cells were transferred to irradiated WT (white) or *Card9*^*-/-*^ (gray) recipients. Another group is *Card9*^*-/-*^ BM transferred to WT recipients (orange). *n*=4-5 mice/group. **(h)** Negative control using NFAT-GFP reporter cells lacking mDectin-1/mCD3ζ. The control cells were stimulated with hk*Mtb* (200 μg/ml) for 20 hrs.

**Supplementary Figure 4.**
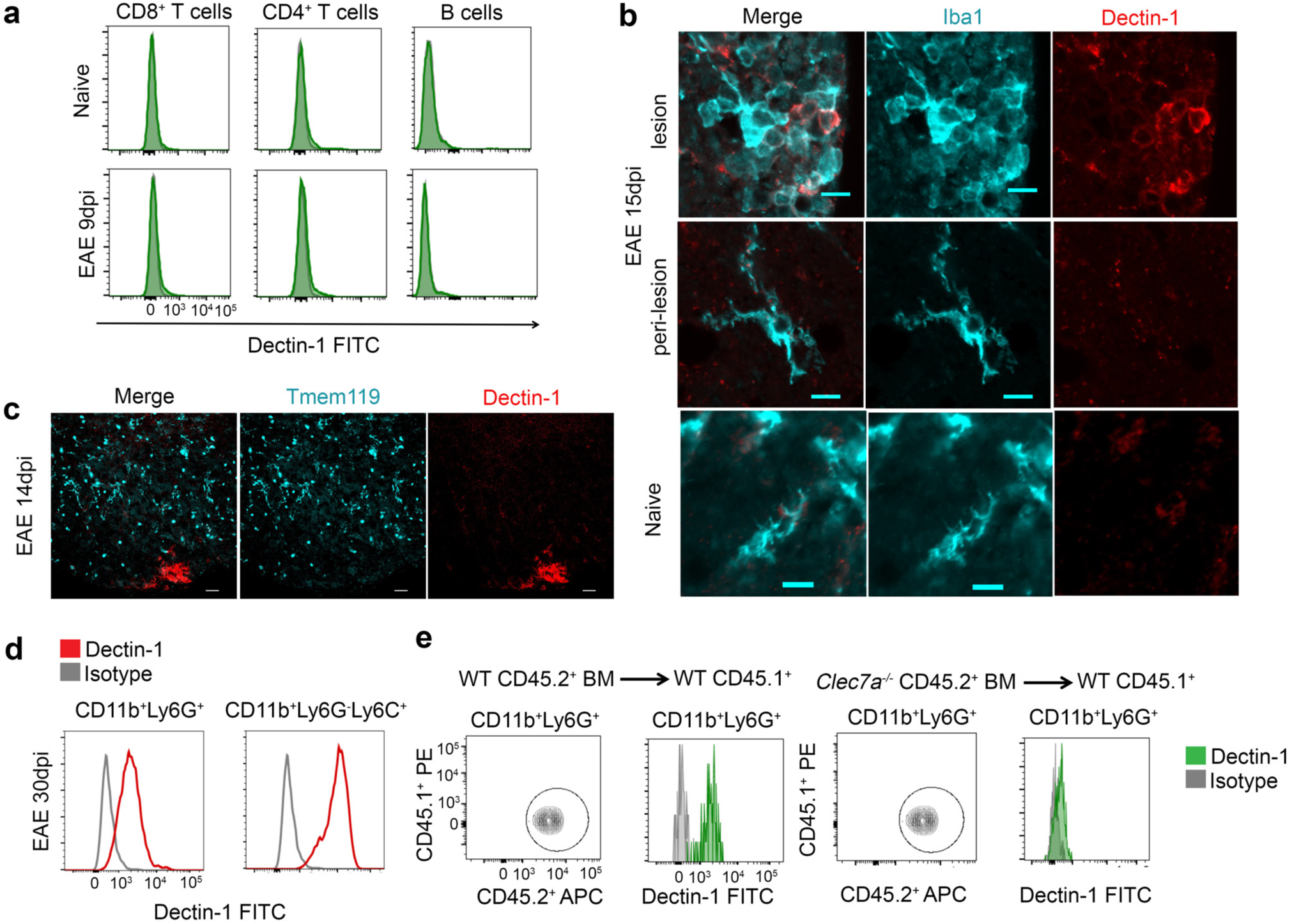
Flow cytometry and immunofluorescent microscopy evaluating Dectin-1 expression. **(a)** Flow cytometry analysis of Dectin-1 expression on splenic lymphocytes in naïve and 9-dpi EAE mice. **(b)** Detection of Dectin-1 expression on Iba1^+^ cells in the lumbar SC white matter (WM), showing lesion and peri-lesion areas, from EAE 15-dpi. Corresponding SC-WM regions from naïve mice also shown. Scale 10μm. **(c)** Detection of Dectin-1 and Tmem119 expression in the lumber SC-WM from mice with EAE 14-dpi. Scale 100μm. Representative of *n*=5 mice/group from 2 independent experiments. **(d)** Dectin-1 (red) expression on SC neutrophils (CD11b^+^Ly6G^+^) and monocytes (CD11b^+^Ly6G^-^Ly6C^+^) at EAE 30-dpi. Histogram of the isotype control is also indicated (grey). Data representative of 2 independent experiments. **(e)** Confirming reconstitution of adoptively transferred BM cells (CD45.2), which were WT or *Clec7a*^*-/-*^, in recipients (CD45.1). Neutrophils (CD11b^+^Ly6G^+^) in peripheral blood of BM chimera were analyzed 8 weeks after BM cell transfer and showed complete chimerism. Data representative of 2 independent experiments.

**Supplementary Figure 5.**
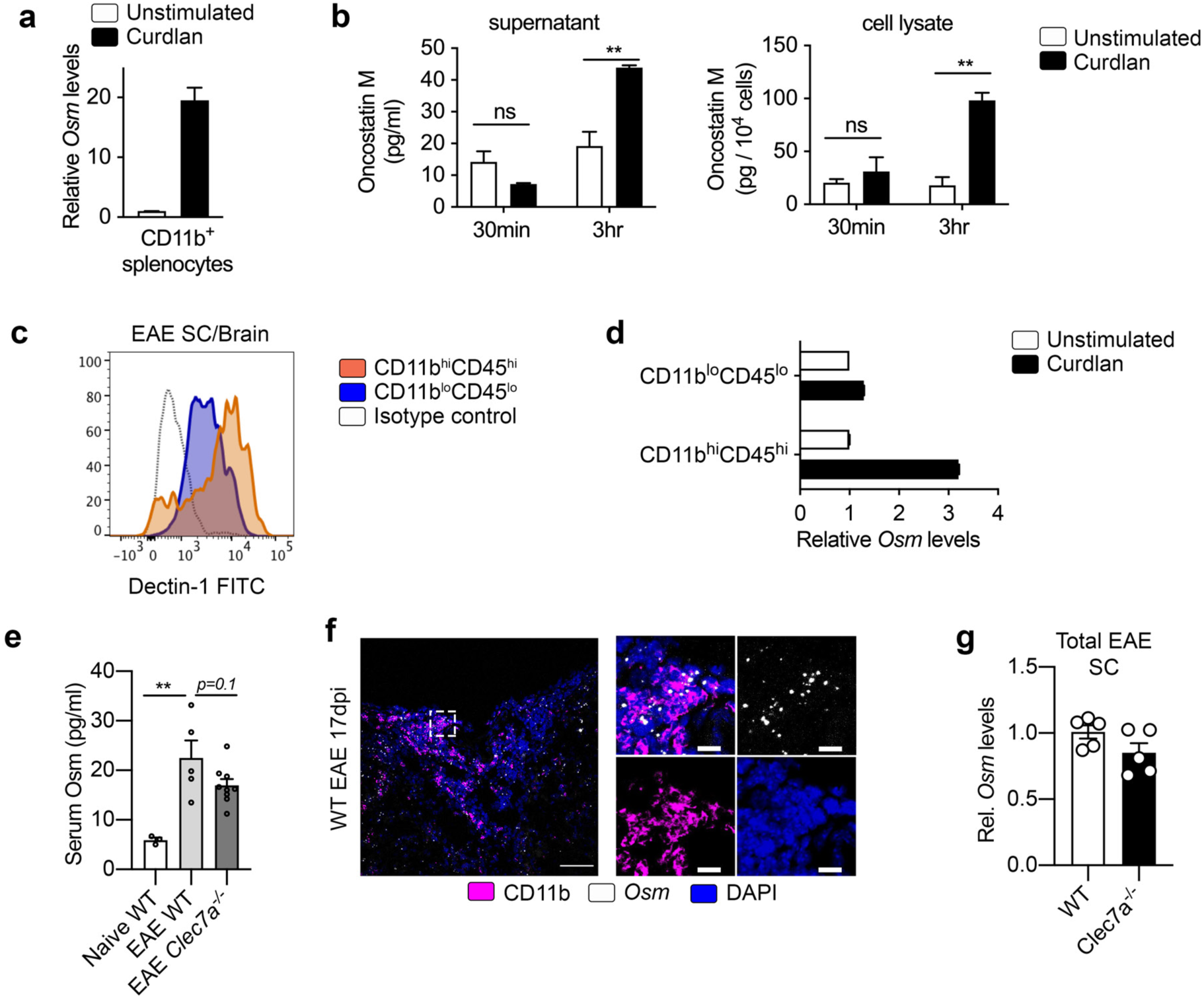
Validation and quantification of Oncostatin M expression. **(a)** *Ex vivo* stimulation of FACS-sorted splenic myeloid cells (CD11b^+^) stimulated with curdlan (100 μg/ml) for 3 hrs. Data representative of two independent experiments. **(b)** Osm protein produced by BM neutrophils, evaluated in tissue culture supernatants and cell lysates at 30 min or 3hrs after curdlan stimulation (100μg/ml). Data representative of 2 independent experiments. **(c, d)** CD11b^lo^CD45^lo^ and CD11b^hi^CD45^hi^ cells FACS-sorted from SC and brain of EAE mice at 30-dpi, evaluated for Dectin-1 expression by flow cytometry (c) and for *Osm* expression by RT-qPCR after 3hr *ex vivo* stimulation with curdlan (100μg/ml) (d). Samples pooled from 2 mice each. Representative of 2 independent experiments. **(e)** Serum Osm protein levels from naïve WT and 9-dpi EAE WT and *Clec7a*^*-/-*^ mice. One data point is from one mouse. Representative of two independent experiments. **(f)** Representative images of *Osm* mRNA (RNAscope ISH) and CD11b protein (IF) in the lumbar SC-WM from WT mice at 17-dpi EAE. Scale 100μm in main panels and 10μm in inset. Representative of 2 independent experiments. **(g)** *Osm* gene expression, determined by RT-qPCR, in total SC homogenates from EAE 14-dpi WT and *Clec7a*^*-/-*^ mice (*n*=5 mice/group, combined from 2 independent experiments).

**Supplementary Figure 6.**
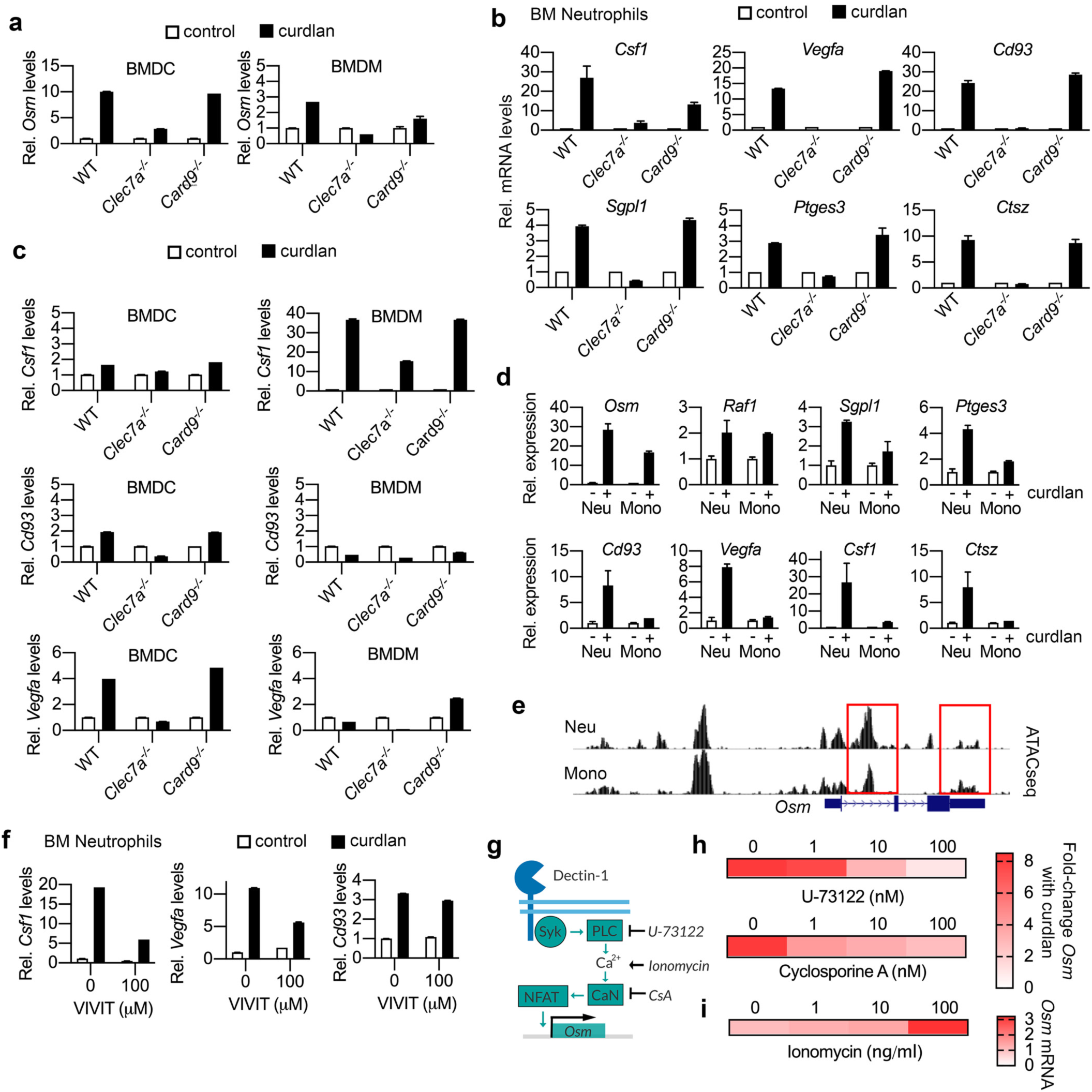
Validation of RNA-seq profiling of Card9-independent transcriptional program by Dectin-1 stimulation. **(a-d)** Levels of mRNA levels were determined by RT-qPCR analysis. *Osm* mRNA levels in BMDCs and BMDMs, stimulated with curdlan (100 μg/ml) for 3 hrs (a). Selected Card9-independent genes in WT, *Clec7a*^*-/-*^, and *Card9*^*-/-*^ BM neutrophils, stimulated with curdlan (100 μg/ml) for 3 hrs (b). Selected Card9-independent genes in WT, *Clec7a*^*-/-*^, and *Card9*^*-/-*^ BMDMs and BMDCs, stimulated with curdlan (100 μg/ml) for 3 hrs (c). RT-qPCR for selected genes in WT BM neutrophils or monocytes stimulated with curdlan (100μg/ml) for 3 hrs (d). **(e)** ATACseq ImmGen data for the *Osm* locus in BM neutrophils or blood Ly6C^+^ monocytes. Boxes indicate open chromatin regions (OCRs) with predicted NFAT binding sites indicated with red rectangles. **(f)** mRNA levels of selected Card9-independent genes in WT BM neutrophils, stimulated with curdlan (100 μg/ml) for 3 hrs after 1 hr pretreatment with an NFAT-inhibitor, VIVIT (100μM). **(g)** Schematic of small molecule targets. **(h, i)** *Osm* mRNA levels in WT BM neutrophils stimulated with curdlan (100 μg/ml) for 3hrs after 1 hr pretreatment indicated small molecules (h), or with ionomycin (i). Data representative of at least 2 independent experiments (a-d, f, h, i).

**Supplementary Figure 7.**
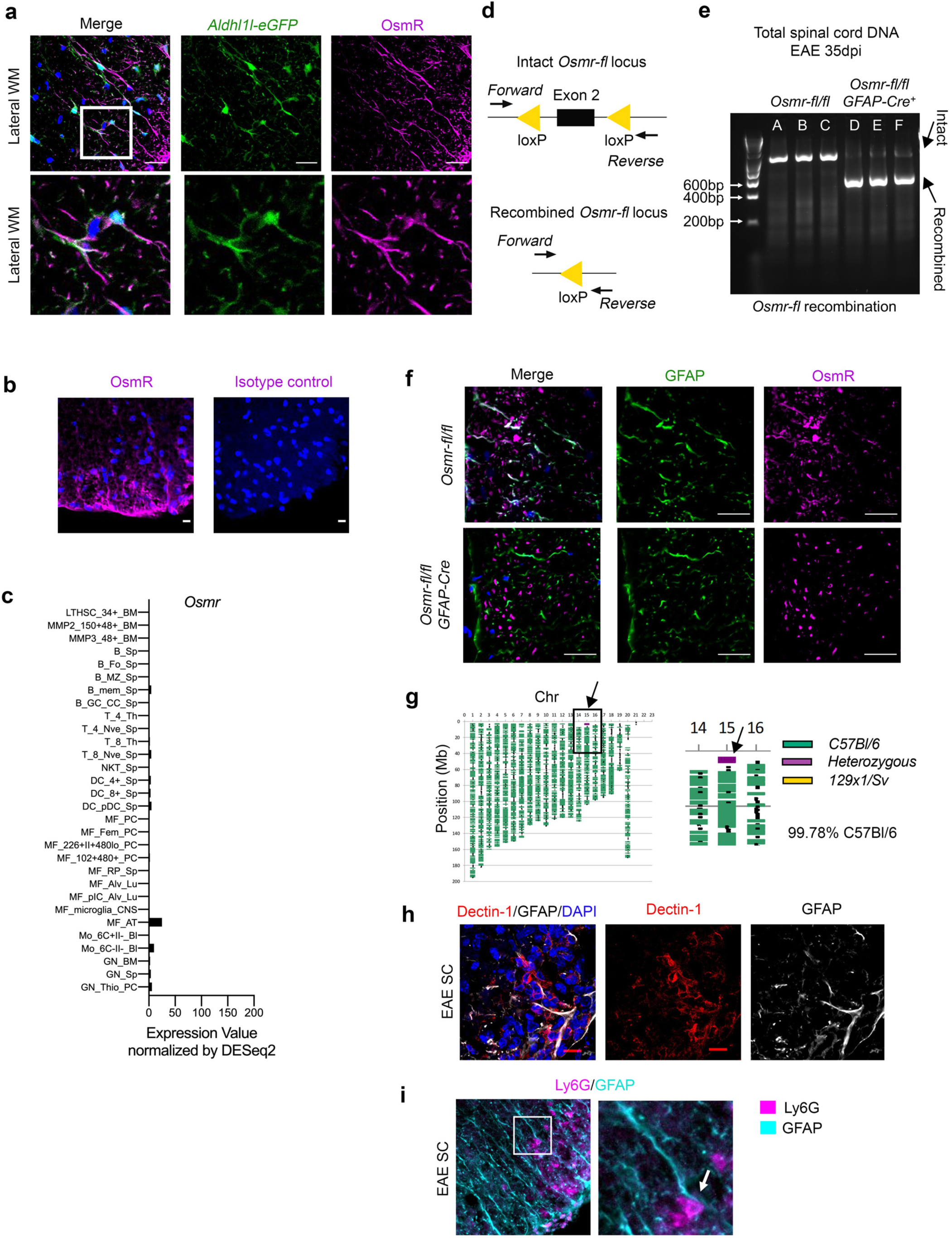
Validation of OsmR expression and targeted deletion in astrocytes. **(a)** Detection of OsmR (magenta) in astrocytes with DAPI staining (blue) in the lumbar SC-WM of naïve *Aldhl1l-eGFP* (green) mouse, scale 25μm. Representative of *n*=3 mice. **(b)** Detection of OsmR (magenta) with DAPI staining (blue) in the lumbar SC-WM of naïve mice. Right panel shows isotype control of OsmR Ab. Scale 10μm. Data representative of *n*=3 mice from 2 independent experiments. **(c)** *Osmr* gene expression in selected immune populations. Data was obtained from Immgen. **(d, e)** Confirming LoxP recombination in *Osmr*^*fl/fl*^;*GFAP*^*Cre*^ mice. Diagram of the *Osmr*^*fl/fl*^ locus with PCR primer annealing positions (d). Results of PCR reaction from total spinal cord DNA (35-dpi EAE) in *Osmr*^*fl/fl*^;*GFAP*^*Cre*^ and *Osmr*^*fl/fl*^ mice using *n=3* mice/group, representative of 2 independent experiments (e). Strong amplification of a recombined DNA amplicon. The remaining long amplicon reflects cells without Cre-recombination. **(f)** OsmR (magenta) expression counterstained for GFAP (green) and DAPI (blue) in lumbar SC (ventral WM) of naïve *Osmr*^*fl/fl*^ and *Osmr*^*fl/fl*^;*GFAP*^*Cre*^ mice. Scale, 25μm. Representative of *n*=5 mice/group combined from 2 independent experiments **(g)** Diagram of the chromosome genetic background of *Osmr*^*fl/+*^ mouse backcrossed to C57BL/6. The purple box includes the *Osmr* locus on Chr 15, 3.30 cM. **(h, i)** Detection of Dectin-1 (red) and GFAP (white) (h) or Ly6G (magenta) and GFAP (cyan) (i) in lumbar SC-WM of EAE 17dpi mice, scale 10μm.

**Supplementary Table 1.**
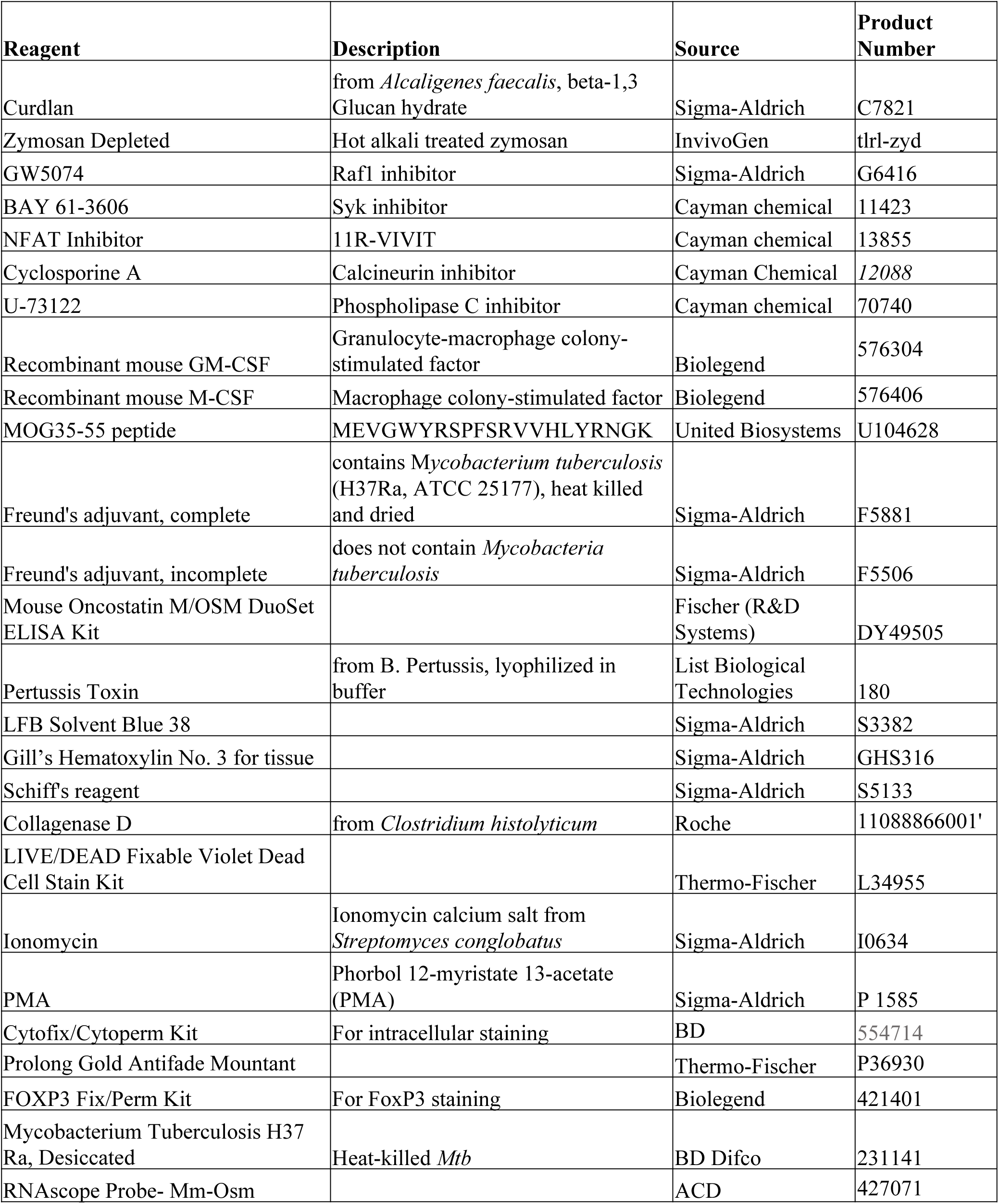
Key reagents.

**Supplementary Table 2.**
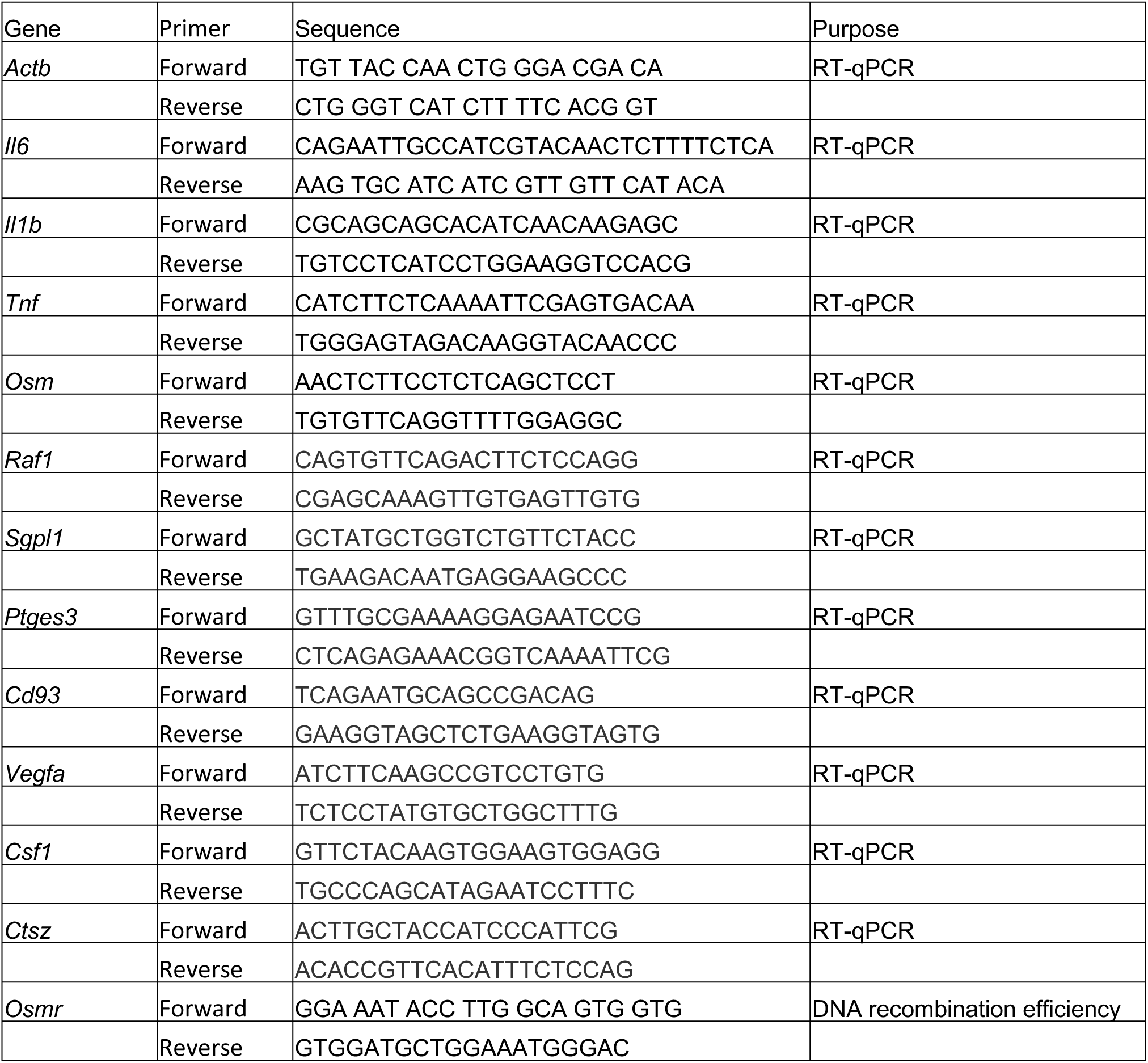
Primer sequences.

